# Inferring antifungal drug synergy from *Candidozyma auris* optical density data using Bayesian mechanistic modelling

**DOI:** 10.64898/2026.06.19.733372

**Authors:** Tara Hameed, Larissa L. H. John, Elaine Bignell, Reiko J. Tanaka

**Author notes:** (R.J.T.).

## Abstract

Antifungal drug-resistant *Candidozyma auris* (*C. auris*) is a threat to human health worldwide. Combination antifungal drug therapy has emerged as a promising approach to combat drug-resistant *C. auris* because some drugs interact synergistically to increase fungal clearance when co-administered. Moreover, combination regimens that either rapidly act or completely kill *C. auris* could mitigate development of on-treatment resistance. However, traditional checkerboard methods to identify synergistic drug combinations only inspect fungal growth at a single timepoint. As a result, they cannot be used to estimate the rate of drug-action or to hypothesise on fungicidal or fungistatic drug-action. Mechanistic modelling would allow us to quantify time-dependent drug-action and infer killing or inhibitory action, but these models are usually fit to direct measurements of fungal growth whose collection is currently not scalable to many time-points and drug combinations. In this paper, we propose a Bayesian mechanistic modelling approach that could detect drug-synergy, estimate drug-action over time and investigate fungicidal or fungistatic drug-activity from optical density (OD_600_) data alone. OD_600_ is quicker and easier to collect than direct measurements of fungal growth and therefore more amenable to high-throughput susceptibility testing. By fitting our model to time-course OD_600_ data of a multi-drug-resistant *C. auris* isolate growing in mono- and combination drug regimens, we successfully inferred synergy between previously confirmed synergistic antifungal drugs (anidulafungin with manogepix or with 5-flucytosine) and linked our model’s inferred kinetic parameters to fungicidal and fungistatic action on *C. auris* growth, which matched drug-activity reported in literature where known. We validated that our model outperformed baseline logistic and Gompertz models using cross validation stratified by OD_600_ replicates. Our results represent the much-needed groundwork for identifying drug combinations for subsequent experimental testing for use in clinics based on their synergy, temporal drug-action and fungicidal or fungistatic activities inferred from OD_600_ data alone.

**Author Summary:** There is an urgent need to locate novel treatments to better treat antifungal drug-resistant *Candidozyma auris* infections. Combination therapy is a promising approach where two or more antifungal drugs are administered and interact synergistically to enhance fungal clearance. If these combinations are fast acting or eradicate fungi through killing, then they could also reduce the chance of resistance developing during treatment. The synergy of antifungal drug combinations is currently assessed by checkerboard methodologies that compare fungal growth under drug combinations to that under a single drug. However, checkerboard methodologies record only one time-point. Hence, they cannot evaluate drug combinations’ timeframe of action and follow-up studies are required to determine which combinations could optimally enhance killing. We developed a Bayesian mechanistic model that could detect synergy between drugs, estimate rates of drug-action and investigate killing and inhibition drug-action using only optical density (OD_600_) data of *C. auris*. OD_600_-based measurement of fungal growth is more amenable to large-scale drug testing than data typically used for mechanistic modelling, such as microscopy data. This work serves as a foundation for more targeted drug testing that identifies promising drug combinations based on their inferred drug-synergy and hypothesised killing (or inhibition) rates.

## Introduction

*Candidozyma auris* (*C. auris*) can cause severe, life-threatening infections in humans [1]. Although antifungal drugs are the standard treatment, *C. auris* has been found to be multi-drug resistant [2], with some strains even demonstrating pan-drug resistance [2,3]. As a result, antifungal drug-resistant *C. auris* has been highlighted as an urgent public health issue by both the World Health Organisation (WHO) [4] and the US Centers for Disease Control and Prevention (CDC) [5].

Combination antifungal drug therapy has recently emerged as a promising approach to combat antifungal drug-resistant *C. auris* infections [6]. It involves administration of two or more antifungal drugs that interact to increase fungal clearance. For example, combinations of antifungal drugs such as micafungin with amphotericin B [7] or flucytosine with other antifungal drugs [8] have been found to act synergistically when co-administered. Such drug synergy is not only promising for treating refractory *C. auris* infections [8] but could also reduce the required dosage of antifungal drugs, thus reducing treatment cost [6]. It has also been suggested to potentially mitigate the emergence of on-treatment resistance [6]. Rapidly effective synergistic drugs have the potential to limit the window of opportunity for resistance emergence and fungicidal drug combinations could eradicate fungi prior to resistance occurring. To identify such synergistic drug combinations, we need to be able to detect drug synergy, identify rapidly effective regimens and fungicidal combinations. Moreover, individual responses of many clinical isolates must be tested in an efficient and high-throughput manner since different *C. auris* isolates have varying responses to different drug combinations [6,9].

Traditionally, checkerboard analyses are typically used to determine which antifungal drugs should be applied to achieve drug synergy for a given isolate [10,11] by evaluating fungal growth at a single time point for a range of drug combinations and concentrations compared to monotherapies. However, standard checkerboard approaches measure fungal growth only at a single time-point, so cannot support quantification of time-dependent (dynamic) antifungal drug-action or hypothesise on fungicidal versus fungistatic drug-activity without further follow-up studies. Quantifying dynamic drug-action is required to calculate and compare rates of action over time and subsequently identify rapid-acting regimens. Some drug-action may even be obfuscated if we look at only snapshot differences in growth; for example, a drug that has little antifungal action over time may have similar end-point readings to a drug that quickly acts but then allows fungal regrowth to occur. Inferring fungicidal or fungistatic activity can accelerate identification of fungicidal drug-combination regimens for a given isolate. Additionally, checkerboard experiments are time- and labour-intensive, making them currently unsuitable for large-scale testing. To rapidly identify which combinations of antifungal drugs could exhibit quick-acting fungicidal regimens during routine susceptibility testing, a more efficient, temporal and mechanistic evaluation of drug-action will be essential.

Time-dependent drug-action can be inferred by mathematical models of population growth, especially mechanistic models that are fit to time-course data of fungi growing in different antifungal drug conditions. Analysis of the estimated model parameter values can infer fungistatic and fungicidal drug-action, which serve as hypotheses for subsequent experimental testing and validation. Once the model is validated, various growth outcomes in drug can be quickly and efficiently explored *in silico*. However, collecting time-course data for model parameter fitting usually requires direct measurement of *C. auris* growing over time, using microscopy methods or recording colony forming units (CFUs). Such measurement is prohibitively time- and labour-intensive to collect for multiple time-points and drug combinations during susceptibility testing.

In this paper, we address these issues by developing a Bayesian mechanistic model to detect antifungal drug synergy, quantify drug-action over time, and link this drug-action to *C. auris* killing and growth-inhibition using only time-course optical density (OD) data. OD is an indirect measure of growth that is quicker and easier to collect than direct measurements and therefore more amenable to large-scale susceptibility testing. Using OD data of a multi-drug-resistant isolate of *C. auris*, we show that our model could infer synergistic drug-combinations and make predictions of potential fungicidal or fungistatic drug-action that can be subsequently experimentally verified.

## Results

### OD_600_ data of C. auris growth

OD data of a multi-drug-resistant B12663 *C. auris* isolate [41] measured at a wavelength of 600 nm (OD_600_) over 48 hours was collected (**Fig 1**). The B12663 *C. auris* isolate was grown in Roswell Park Memorial Institute (RPMI) media alone or co-incubated with five antifungal drug regimens: 5-flucytosine (5FC), anidulafungin (AFG), manogepix (MGX) or a combination of AFG with 5FC (AFG+5FC) or AFG with MGX (AFG+MGX) at each drug’s minimum inhibitory concentration (MIC50), the concentration required to reduce growth by 50%, which were previously determined in [9]. Combinations with AFG were prioritised because AFG is an echinocandin, which is the current recommended first line antifungal drug [12,13], almost always used in monotherapy and against which resistance is steadily emerging [14].

**Fig 1.**
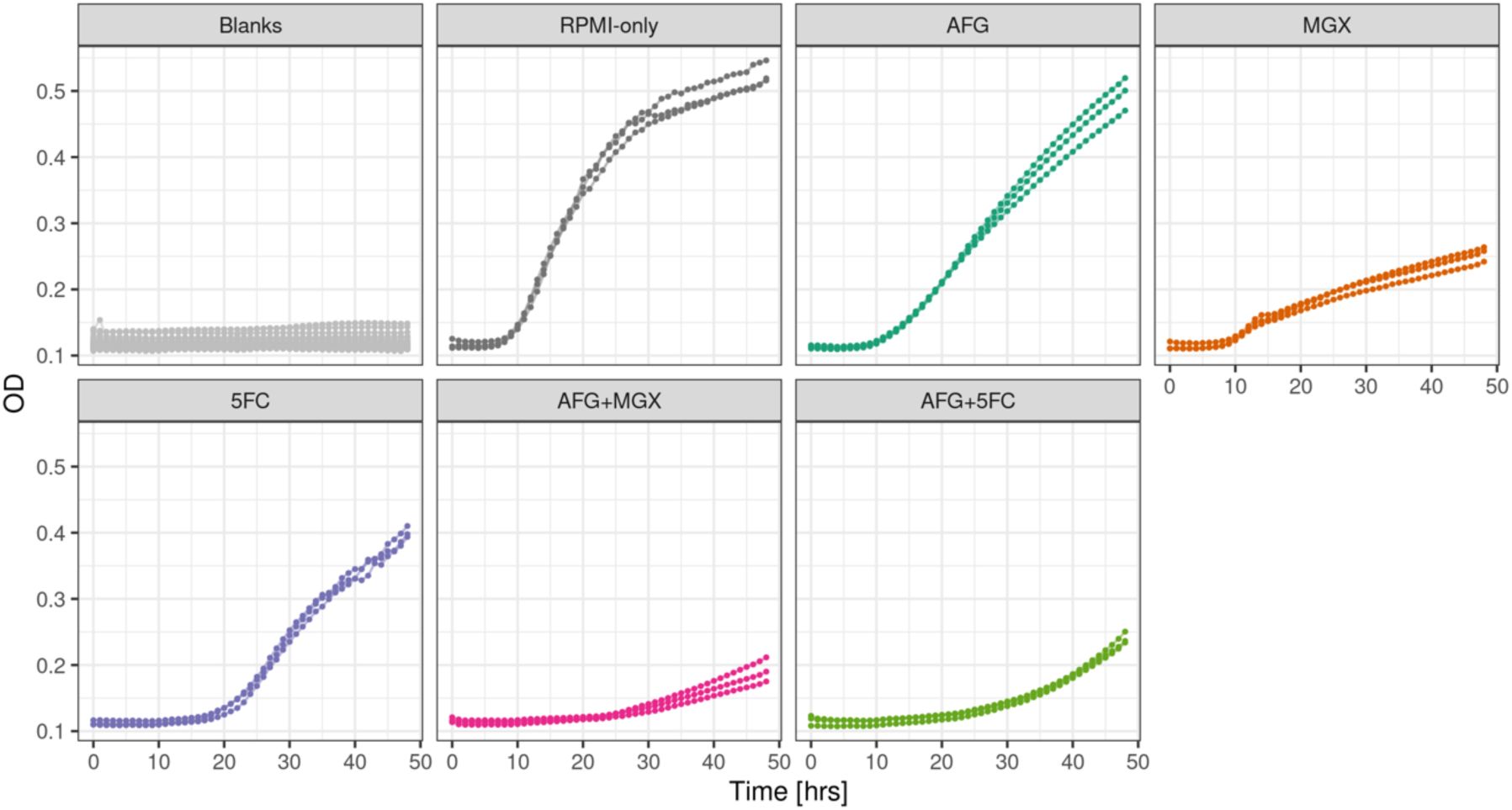
Experimental OD_600_ data. OD_600_ data (*dots*) of wells with RPMI and H_2_0 (Blanks, *light grey*) or B12663 *C. auris* growing over 48 hours in RPMI (*dark grey*) with 8 [mg/L] AFG (*dark green*), 0.03 [mg/L] MGX (*orange*), 0.25 [mg/L] 5FC (*purple*) or their combinations (AFG+MGX (*pink*) or AFG+5FC (*light green*). OD_600_ replicates from the same well are joined by a line.

### A Bayesian mechanistic model of C. auris growing in drug-free conditions

We first proposed and evaluated models for OD_600_ data of B12663 *C. auris* growing in only drug-free conditions (**Fig 1**, *Grey*). To account for OD being an indirect measurement, we modelled the OD_600_ data of the B12663 isolate as distributed around a linear transform of latent fungal growth [15]. We considered four population growth ordinary differential equation (ODE) models for the latent fungal growth: exponential, logistic [16], Gompertz [17] or Edwards [18,19] (see **Methods,** *Mechanistic models*).

We assessed all models’ predictive performance by cross validation (CV) stratified by OD_600_ replicates (**Fig 2**, *Drug Free*). The Gompertz and Edwards models achieved the highest predictive performance compared to the exponential and logistic models, as demonstrated by their lower relative log predictive density (LPD) and root mean squared error (RMSE) on held-out B12663 OD_600_ replicates. They also had the best fit to the drug-free OD_600_ data, indicated by the lowest RMSE on the training data over all the folds (**Fig S1**, *Drug Free*). The predictive performance was worse for models that did not account for OD being an indirect measure (Gompertz and Edwards (OD direct) models) (**Fig 2**, *Drug Free*).

**Fig 2.**
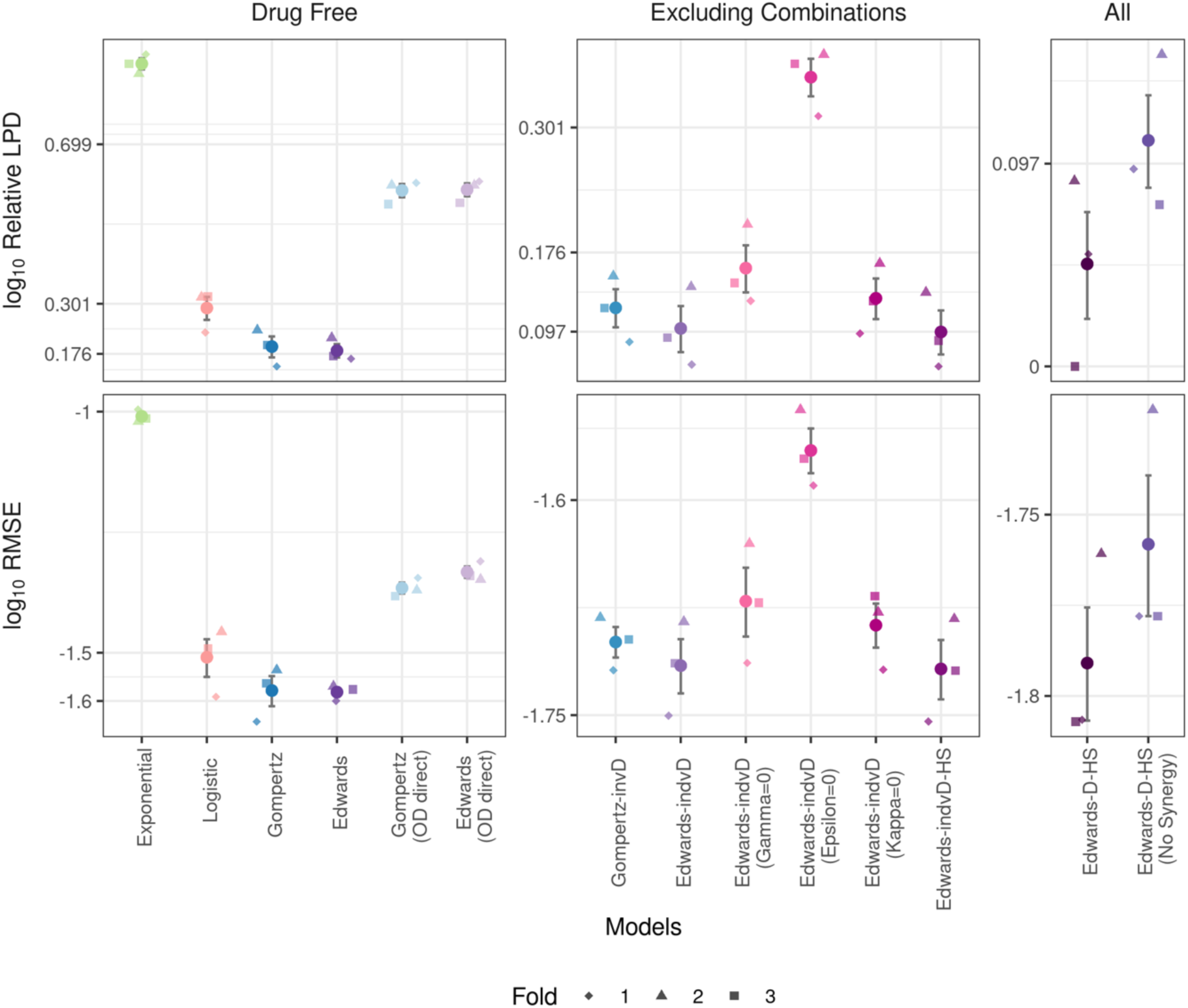
Cross validation with OD_600_ data of B12663 *C. auris* Mean relative LPD ((maximum LPD)-LPD + 1) and mean RMSE (three significant figures (3.s.f)) per testing OD_600_ replicate averaged over testing replicates for each fold (*k* = 3). Mean and standard errors over folds are shown as dots with grey error bars. Models were assessed on drug-free OD_600_ data (*left*), with the monotherapeutic regimen OD_600_ data (*centre*) and with combination drug regimen data (*right*). Lower values indicate better predictive performance for both metrics.

### Extending the best performing models in drug-free conditions to include drug-action

To allow for inference of fungicidal or fungistatic drug-action over time, we extended the Gompertz and Edwards models, the best performing models on drug-free data, to include individual antifungal drug-action during monotherapy (see **Methods,** *Mechanistic models*). In both models, we added a direct-killing term to allow for inference of killing rates and included drug-induced inhibition of growth dynamics. The resulting Gompertz-indvD and Edwards-indvD models included three mechanisms for individual drug-action of each drug *j* ∈ {AFG, MGX, 5FC}: (1) direct killing at rate *k*_j_, (2) growth rate inhibition at rate γ_j_, and (3) reduction of the carrying capacity in the Gompertz-indvD model or enhancement of *C. auris’* rate of growth-impedance due to population size in the Edwards-indvD model, at rate ε_*j*_, where a parameter that controls growth-impedance due to population size was already present in the drug-free Edwards model. Whilst direct killing, *k*_j_, and inhibition of the growth rate, γ_j_, represent direct drug-action, enhancement of the growth-impedance, ε_j_, represents a biologically mediated response to drug.

The Edwards-indvD model had improved fitting and predictive performance compared to the Gompertz-indvD (**Fig 2** and **Fig S1,** *Excluding Combinations*), where it scored lower relative LPD and RMSE in each fold. We retained all three drug-action mechanisms in the Edwards-indvD model because removing any of the mechanisms (by setting γ_j_, _j_, *k*_j_ = 0) worsened the CV predictive performance across all folds. We further placed a regularised horseshoe (HS) prior [20] on the drug-action parameters (Edwards-indvD-HS model) to prevent overfitting, where regularisation prevents all three drug-action mechanisms being inferred for a single drug if all are not needed. We confirmed that the Edwards-indvD-HS model did not have worse CV performance and so decided to use the Edwards-indvD-HS model below.

To fit to all data, including combination drug-conditions, we extended the Edwards-indvD-HS model to include additive drug-action by summing individual drug-action terms in the ODE that correspond to a given drug combination, for example adding AFG and MGX terms in the model for AFG+MGX drug-action. Moreover, an extra interaction term was added per drug combination to infer any positive interactions (synergy) between drugs (see **Methods,** *Mechanistic models*). The resulting Edwards-D-HS model contains parameters, γ_j_, ε_j_and *k*_j_, that quantify the strengths of antifungal drug-action for *j* ∈ {AFG, MGX, 5FC} and drug-synergy for *j* ∈ {AFG: MGX, AFG: 5FC}, which were inferred during fitting.

The Edwards-D-HS model achieved the highest predictive and fitting performance on testing (**Fig 2**, *All*) and training (**Fig S1**, *All*) data, respectively, on average over the folds compared to the other models, as evaluated by CV stratified by OD_600_ replicates of *C. auris* growing in RPMI and all drug conditions. The predictive performance during CV was worsened for the Edwards-D-HS (No Synergy) model that had no synergy, implemented by setting the added interaction parameter to zero, and only additive drug-action for the combinations, suggesting the importance of the synergy terms in the Edwards-D-HS model.

Since the Gompertz-indvD model had overlapping relative LPD and RMSE standard errors with the Edwards-indvD (**Fig 2**, *Excluding Combinations*), we also extended the Gompertz-indvD to include combination drug-conditions and confirmed that the resultant Gompertz-D-HS model had worse fitting and predictive performance than the Edwards-D-HS on average for both training and test data (**Fig S1**, *All*). Moreover, removing synergy from the Gompertz-D-HS also worsened predictive performance, suggesting the synergy terms were also important for the Gompertz-based models. We additionally confirmed that there was no clear difference in the identifiability of drug-action parameters between the Gompertz- and Edwards-D-HS models, measured by the drug-action parameter’s credible interval coverage and posterior contraction (*s*) and *z*-scores [21] (**S1 Text**, *Section B* and **Fig S2**). We did not extend the Edwards-indvD (*k*_j_= 0) to include drug combinations because inferring the killing rate, *k*_j_, is key to inferring fungicidal action and setting *k*_j_ = 0 reduced CV performance in each fold. We instead verified the identifiability of all drug-action parameters in the Edwards-D-HS model using a fake data check (**S1 Text,** *Section B* and **Fig S3**).

### Detection of drug synergy

The Edwards-D-HS model was successfully fit to all the OD_600_ data, as confirmed by posterior predictive checks (**Fig 3a**). For each of the inferred drug-action parameters, γ_j_, ε_j_and *k*_j_ for *j* ∈ {AFG, MGX, 5FC, AFG: MGX, AFG: 5FC}, we checked the corresponding inferred fungicidal or fungistatic drug-action against known drug-activity in literature. The inferred parameters were considered as non-zero if the lower bound of their 95% credible interval was above 10^-4^. A perturbation of 10^-4^ from 0 for the drug-action parameters had no visible change to model output, resulted in less than 0.1% change in the ODE solution of viable growth and the difference in model output from drug-free simulations had an RMSE that was at least an order of magnitude below the lowest RMSE recorded during CV on testing or training data (**S1 Text**, *Section C* and **Fig S4**).

**Fig 3.**
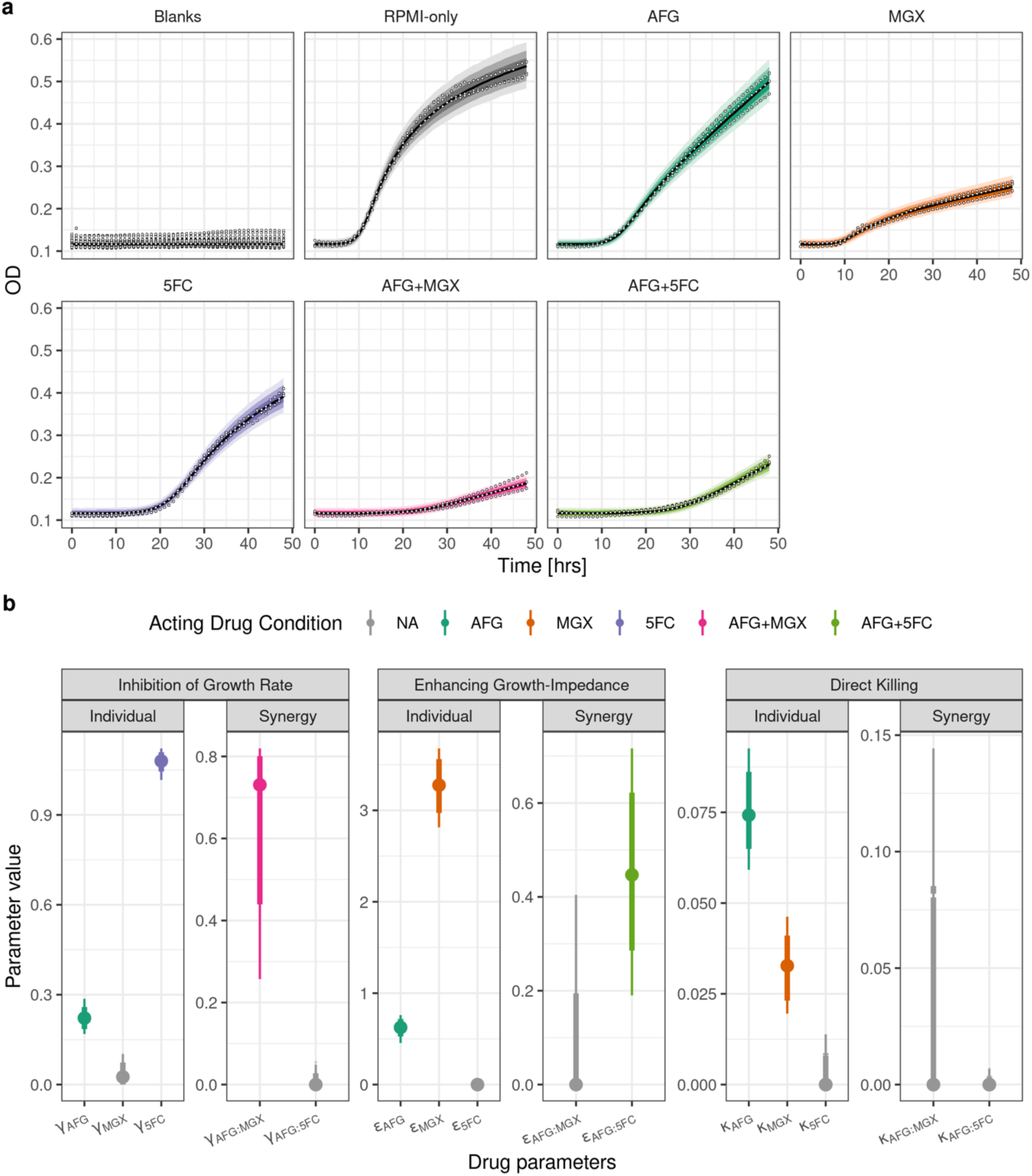
Edwards-D-HS model fit to OD_600_ data and inferred parameter values. (**a**) Posterior predictive distribution (solid lines and shades are medians and 95%, 80%, 50% highest density credible intervals, respectively). OD_600_ data (*white dots*) of B12663 *C. auris* grown in the presence of of AFG, 5FC or MGX, or two-way combinations thereof, over time. (**b**) Estimated parameter values (median (*dots*), 95% (*thin line*) and 80% (*thick line*) highest density credible intervals) for AFG (*green*), MGX (*orange*) and 5FC (*purple*) individual drug-action and AFG+MGX (*pink*) and AFG+5FC (*light green*) drug synergy through inhibiting the growth rate (γ.), enhancing *C. auris’* growth-impedance (ε.), and direct killing (*k*.) (*left to right*). Parameter values where inferred 95% credible intervals excluded 10^-4^ were considered to be non-zero and those that included 10^-4^ are shaded grey.

For AFG, our model inferred non-zero values for γ_AFG_, ε_AFG_ and *k*_AFG_ (**Fig 3b**, *dark green*), indicating that AFG acted to inhibit *C. auris’* growth rate, enhance *C. auris’* growth-impedance and kill *C. auris* in the model. These outcomes are concordant with those of prior *in vitro* experiments that show AFG inhibits *C. auris* growth (fungistatic) and can kill *C. auris* but not enough to be classed as a fungicidal drug [22,23].

For MGX, our model inferred non-zero parameter values for ε_MGX_ and *k*_MGX_ (**Fig 3b**, *orange*), corresponding to the enhancement rate of *C. auris’* growth-impedance and direct killing rate of *C. auris*. *In vitro* experiments have shown that MGX does have potent activity against *C. auris* [24–26]. However, no studies have confirmed if MGX’s main activity against *C. auris* is killing or inhibition-based yet, as far as we are aware. In general, MGX is known to inhibit the Gwt1 enzyme, which is crucial for fungal cell wall integrity [27], causing growth defects, such as loss of the ability to form biofilms [27] or filament [28]. This has been documented to cause some loss of viability for fungi [29] but, generally, MGX is considered to inhibit fungal growth [30,31].

5FC is documented as having a fungistatic activity against *C. auris* [32]. Likewise, our model inferred the growth rate inhibition parameter γ_5FC_as the only non-zero drug-action parameter for 5FC (**Fig 3b**, *purple*), whereas the direct killing rate *k*_5FC_had inferred credible intervals whose lower bound was less than 10^-4^.

Finally, our model inferred that both AFG+MGX and AFG+5FC combinations exhibited drug synergy, since inferred credible intervals for γ_AFG:MGX_ and ε_AFG:MGX_ excluded values of 10^-4^ and below (**Fig 3b**, *pink and light green*). Synergistic activity of both drug pairings has previously been revealed *in vitro* [9]. Traditional end-point measurement of fungal growth via checkerboard methodology demonstrated that AFG+MGX and AFG+5FC acted synergistically, as measured by different synergy metrics, and the combinations’ potency was further confirmed by microscopy experiments [9].

The Gompertz-D-HS fit to all the OD_600_ data also inferred γ_AFG_, ε_AFG_, *k*_AFG_, ε_MGX_, *k*_MGX_, γ_5FC_ and γ_AFG:MGX_ to be non-zero for the AFG, MGX, 5FC and AFG+MGX drug conditions (**Fig S5**) and hence also inferred that AFG and MGX exhibited synergy when co-administered. However, the Gompertz-D-HS model failed to robustly infer synergy for the AFG+5FC combination, where all γ_AFG:MGX_, ε_AFG:5FC_ and *k*_AFG:5FC_ had 95% credible intervals whose lower bound included 10^-4^. As the synergy of both AFG+MGX and AFG+5FC has been previously experimentally confirmed [9], this supported the use of our Edwards-D-HS model over the Gompertz-D-HS.

## Discussion

Combination antifungal drug therapy has the potential to combat lethal, drug-resistant *C. auris* infections and additionally limit drug resistance, protecting the durability of life-saving antifungal agents [6]. Detection of synergistic drug combinations for an isolate currently relies on traditional checkerboard methods, which infer synergy using only fungal growth at snapshots in time and therefore cannot be used to identify fast-acting drug regimens or investigate fungicidal or -static activity during routine susceptibility testing. While ODE-based mechanistic models can infer kinetic rates of drug-action and link these rates to mechanisms acting on fungal growth, such as killing or growth-inhibition, the models are usually fit to time-course CFU data that is prohibitively time- and labour-intensive to collect for many drugs, drug-combinations and isolates.

In this paper, we developed a Bayesian mechanistic model of *C. auris* growing in antifungal drug combinations using OD_600_ data (**Fig 1**), which is easier to collect than time-course CFU data, and demonstrated that our model outperformed alternative models during CV (**Fig 2**). By fitting our model to time-course OD_600_ data, we could detect synergy between antifungal drugs already confirmed to act synergistically in literature [9] (**Fig 3**). Moreover, our fitted model estimated kinetic rates of drug-action and linked drug-action to fungicidal and fungistatic action on *C. auris* growth. These fungicidal and fungistatic drug-activities matched existing knowledge of drug-action on *C. auris* in the literature, where known, and can serve as testable hypotheses of drug-activity against B12663 *C. auris* for future experimental validation.

The strength of our Bayesian mechanistic approach is that it allows for inference of growth-inhibition or killing rates that can be experimentally validated, which cannot be inferred by traditional checkerboard-based methods. Once the model’s inferred drug-action are validated in future experiments, the model could be used to explore various growth outcomes in drug, such as varying time of MGX or 5FC administration during AFG combination therapy, through quick and exhaustive simulation *in silico*. Using mechanistic models in this way has the potential to reduce required experiments and accelerate location of rapid-acting and effective drug regimens for future use in clinics. Moreover, our method could also support future drug discovery research by filtering for potentially quick-acting fungicidal combinations during high-throughput drug-screening.

Previous computational models that used OD data to assess antimicrobial efficacy usually evaluated growth at a single endpoint only [33,34], and thus cannot hypothesise on fungistatic or fungicidal action or infer drug-action over time. Some mechanistic models calibrated to time-course data were proposed to characterise dynamic antimicrobial susceptibility of bacteria [35–38] and fungi [39]. However, these models were applied to time-course CFU data (time-kill data) rather than OD, and so are not amenable to high-throughput testing. A mechanistic model fit to time-course OD_600_ was proposed to assess antibiotic efficacy [40], but not used to detect drug-synergy. Moreover, our model additionally accounts for OD being an indirect measure, which was found to be important when modelling OD [15], regularises the drug-action parameters to avoid overfitting, and uses Bayesian inference to estimate the uncertainty of parameter estimates, which is desired when making medical decisions on drug treatments.

A limitation of our proposed model is that it can currently fit only to a single drug concentration for each drug. Future work could look to expand our model by including drug concentration into the model structure, following other previous mechanistic modelling of antimicrobial dose-response modelling [35–39]. Moreover, our model was only tested and fit to a single isolate of *C. auris*. The model would need to be re-fitted and re-evaluated on time-course OD_600_ data of alternative *C. auris* isolates for generalisability. Future work could also look to expand our model to include data from different isolates using a hierarchical structure, which would enable predictions of a new isolate growing in drug.

In summary, we developed a Bayesian mechanistic model that was able to detect antifungal drug synergy, infer time-dependent drug-action and investigate killing or inhibition drug-action for a multi-drug-resistant *C. auris* isolate using only OD_600_ data. This work represents the much-needed foundation for inferring synergistic drug-combinations, their rates of action and identifying experimentally testable killing or inhibition drug-activity, thereby allowing for more targeted susceptibility testing, using only easy-to-collect OD_600_.

## Methods

### OD_600_ Data

B12663 [41] *C. auris* was grown in a 96-well plate over a period of 48 hours and the OD_600_ was recorded. An initial inoculum of 2.5 x 10^2^ cells/μL was used and OD_600_ readings were taken every hour. Inoculum, antifungal drugs and plate were prepared according to EUCAST guidelines [42]. The isolate was grown in RPMI medium alone or with 8 [mg/L] AFG, 0.25 [mg/L] 5FC, 0.03 [mg/L] MGX, 8 [mg/L] AFG + 0.25 [mg/L] 5FC or 8 [mg/L] AFG + 0.03 [mg/L] MGX. The RPMI medium was supplemented with glucose to 2% and buffered at pH 7 using 3-(N-morpholino) propanesulfonic acid (MOPS) at a final concentration of 0.165 mol/L as described in the EUCAST guidelines [42].

### Mechanistic models

We considered 17 mechanistic models (**Table 1**) with varied functions for *C. auris* growth and drug-action when included. Priors for all the models are listed in the Supplementary (**Section A**). No models assumed a delay to the start of growth due to a lack of evidence that *C. auris* exhibits such a delay. The Edwards model was proposed for general microbial kinetics [19] and later adapted for fitting to fungal growth data [18]. It assumes that a population will limit its own growth, like a Gompertz or logistic, but the Edwards model assumes an exponential form for the growth-impedance due to population size.

**Table 1:**
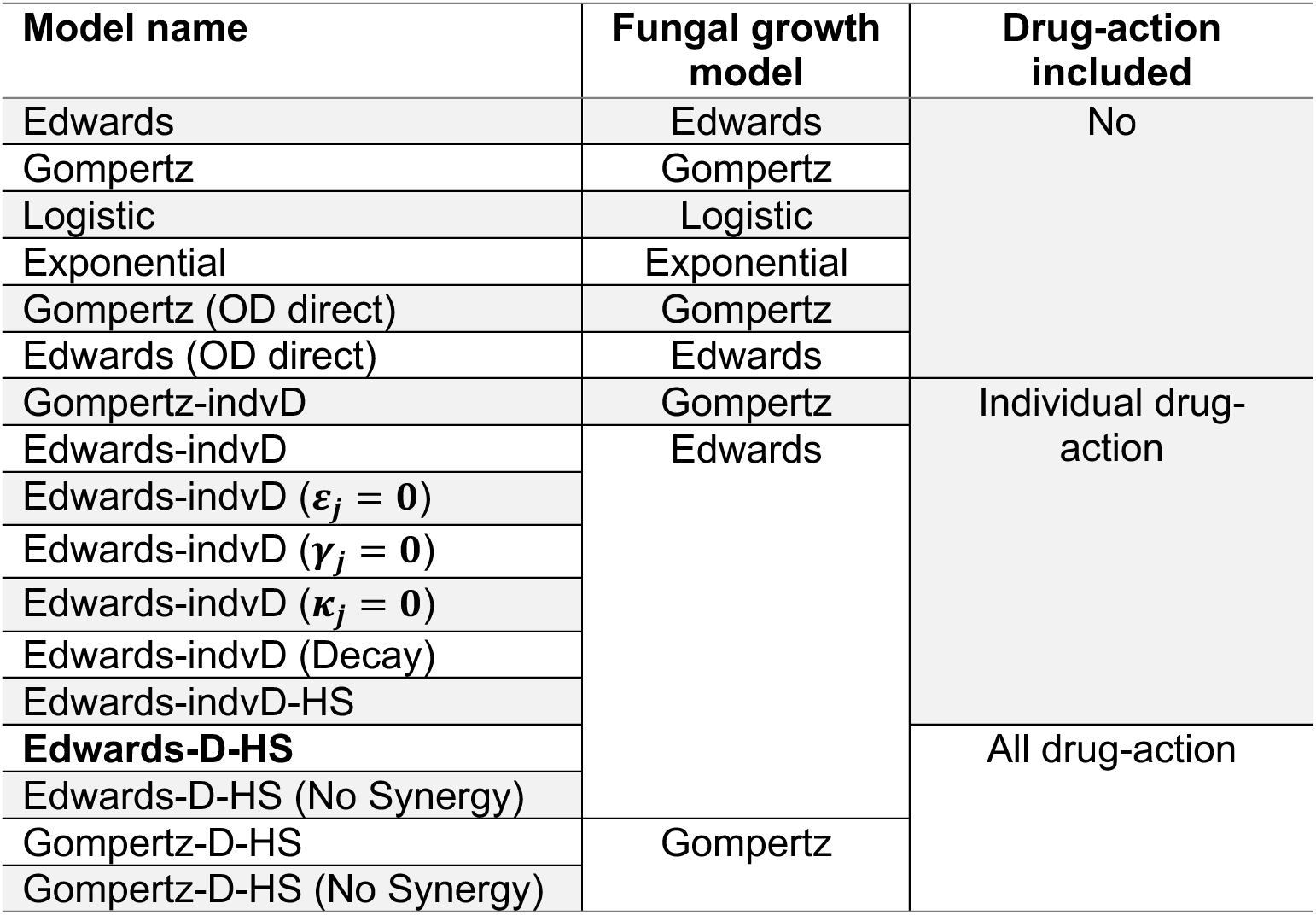
Bayesian mechanistic models considered in this study. Models assumed different functions for *C. auris* growth and included different levels of antifungal drug-action. The Edwards-D-HS model (*bold*) achieved the best predictive performance on average out of all the models.

### Fungal growth models

We considered four models for *C. auris* growth, *f*(*t*), in RMPI only:

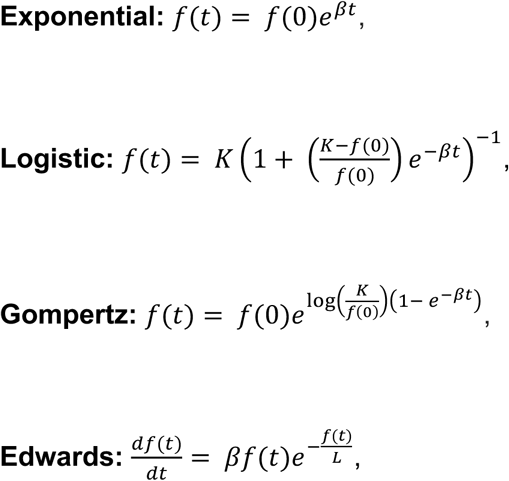

where β [hour^-1^] is the growth rate, *f*(0) [cells/μl] the initial inoculum of *C. auris*, *K* [cells/μl] the carrying capacity, and 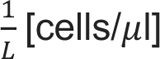 the strength of the *C. auris’* growth-impedance.

The four models for *C. auris* growth were fit to drug-free OD_600_ data following the previously published method to infer fungal growth rates from OD_600_ data [15]. In short, we assumed that the measured OD_600_, *y*_t_, followed a lognormal distribution whose median is the linear transform of *C. auris* growth,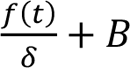:

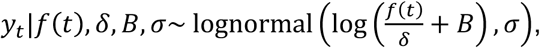

where *B* and δ are the offset and scaling parameters in the linear transform, respectively, and σ is the (multiplicative) noise scale. For the RPMI only wells (blanks), the OD_600_ data is assumed to be distributed as *y*_t_| *B*, σ∼ lognormal(log(*B*), σ).

In the Edwards (OD direct) and Gompertz (OD direct) models, where OD is not assumed to be an indirect measure of growth, the median of the lognormal distribution was set to be *f*(*t*): *y*_t_|*f*(*t*), σ∼ lognormal(log(*f*(*t*)),σ).

### Modelling antifungal drug-action

We proposed the Edwards-indvD and Gompertz-indvD models by including the following three modes of antifungal drug-action. For drug conditions *i* ∈ {No drug, AFG, MGX, 5FC}:

1. Inhibition of the growth rate represented by 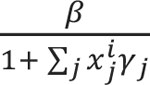, with the inhibition strengths γ_j_ for each of the drugs *j* ∈ {AFG, MGX, 5FC}.
2. Enhancement of *C. auris’* growth-impedance due to population size represented by 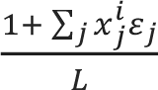 for the Edwards-indvD model or decrease of the carrying-capacity by 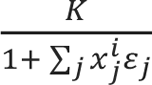 for the Gompertz-indvD model, both with strengths ε_j_.
3. Direct killing of *C. auris* represented by an additional term, 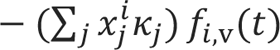, in the growth model with a killing rate, *k*_j_ [hour^-1^], and the growth function, *f*_i,v_(*t*), for viable fungi.

*x^i^_j_ = {0,1}* represents whether the *j*-th drug (*j* ∈ {AFG, MGX, 5FC}) was administered (*x^i^_j_* = 1) or not (*x^i^_j_* = 0) during drug condition *i* (*i* ∈ {No drug, AFG, MGX, 5FC}). The Edwards- and Gompertz-indvD models describe the latent *C. auris* growth function by *f*_i_(*t*) = *f*_i,v_(*t*) + *f*_i,d_(*t*), a sum of the viable (*f*_i,v_(*t*)) and dead (*f*_i,d_(*t*)) *C. auris* for each drug condition *i* because OD readers cannot differentiate between viable and dead fungi. The functions *f*_i,v_(*t*) and *f*_i,d_(*t*) satisfy 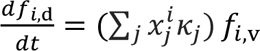 and

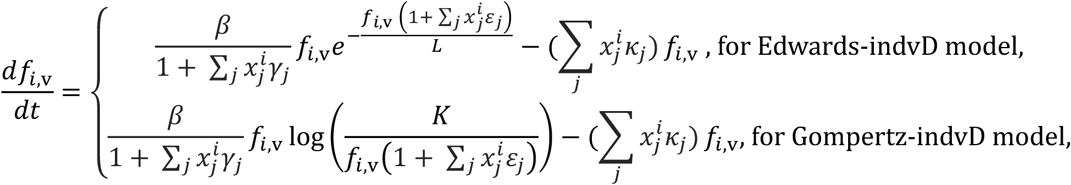

where we use standard linear models for each drug *j, Σ_j_x^i^_j_γ_j_, Σ_j_x^i^_j_γ_ε_* and *Σ_j_x^i^_j_k_j_*, in each of the included drug-action terms. The γ_j_, ε_j_ and *k*_j_ parameter values are bounded to be strictly positive to assume positive drug-actions. These parameters, γ_j_, ε_j_ and *k*_j_, were inferred for all antifungal drugs simultaneously, *j* ∈ {AFG, MGX, 5FC}, to quantify individual drug-action. If their 95% credible intervals had lower bounds above 10^-4^, we concluded that the model estimated non-zero values for those parameters suggesting that the drug *j* acted individually through their corresponding drug-terms acting on *C. auris* growth: inhibition growth, enhancement of growth-impedance or direct killing, for γ_j_, ε_j_ or *k*_j_, respectively. The threshold value of 10^-4^ was selected by a perturbation analysis where we perturbed each of the drug parameters γ_j_, ε_j_ and *k*_j_ by 10^-p^ for *p* ∈ {1, …, 6} from 0 and found *p* ≥ 4 resulted in no visual difference and minimal percentage and RMSE difference in model output (**S1 Text,** *Section C* and **Fig S4**).

Before solving the ODEs, 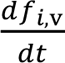 and 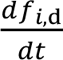 were transformed to be dimensionless in time by scaling *t* by the maximum observed time, *t*_max_, such that the new scaled time 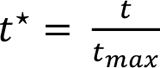 lies in [0, 1] (details in **S1 Text,** *Section A*). Hence, the transformed killing parameters *k^*^_j_ = t_max_ K_j_* are unitless for all *j* ∈ {AFG, MGX, 5FC}, where γ_j_ and ε_j_ were already unitless. This allows all the drug action parameters and their priors to be on the same scale and reduces the computational time taken to solve the ODEs, where the scaled ODEs 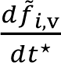 and 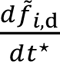 were solved by Stan’s rk45 ODE solver.

The Edwards-indvD (ε_j_ = 0), Edwards-indvD (γ_j_ = 0) and Edwards-indvD (*k*_j_ = 0) models were obtained by setting ε_j_, γ_j_ or *k*_j_ to 0, respectively, for all drugs *j* ∈ {AFG, MGX, 5FC}. Including a decay term for dead fungi (the Edwards-invdD (Decay) model) did not improve the predictive performance during CV compared to the Edwards-indvD model (**Fig S1**, *Excluding Combinations*) and has an extra parameter and hence was not extended to the drug combination data. The Edwards-indvD-HS model has a regularised HS prior [43] on all ε_j_, γ_j_ and *k*_j_ (**S1 Text,** *Section A* for further details).

The Edwards-D-HS model has the same equations as the Edwards-indvD-HS model,

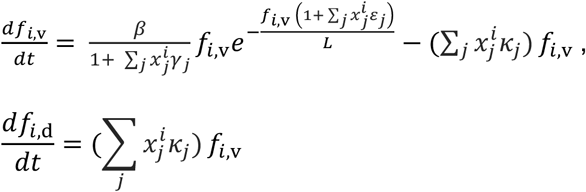

But now *i* indexes all drug conditions in the data, including combinations, *i* ∈ {No drug, AFG, MGX, 5FC, AFG + MGX, AFG + 5FC} and *j* indexes all drugs (AFG, MGX, 5FC) and drug-interactions (AFG:MGX, AFG:5FC), *j* ∈ {AFG, MGX, 5FC, AFG: MGX, AFG: 5FC}. The binary indicator, *x^i^_j_*, is equal to 1 when the *j*-th drug or drug-interaction is present during drug condition *i*. For example, when fitting to data from the drug condition *i* = AFG+MGX, *x^i^_j_ = 1* for *j* ∈ {AFG, MGX, AFG: MGX}, and 0 otherwise, and the drug-action terms in the ODE are ε_AFG_ + ε_MGX_ + ε_AFG:MGX_, γ_AFG_ + γ_MGX_ + γ_AFG:MGX_ and *k*_AFG_ + *k*_MGX_ + *k*_AFG:MGX_, respectively. Hence, in a given drug combination, the model assumed additive drug-action between the individual drugs, ε_j_, γ_j_ and *k*_j_, *j* ∈ {AFG, MGX, 5FC}, and included an extra interaction term per drug combination, ε_j_, γ_j_ and *k*_j_, *j* ∈ {AFG: MGX, AFG: 5FC}, and ε_j_, γ_j_ and *k*_j_ for all *j* were inferred simultaneously during fitting.

If any of ε_j_, γ_j_and *k*_j_ for *j* ∈ {AFG: MGX, AFG: 5FC} were estimated to be positive non-zero values, which we defined as 95% credible interval lower bound > 10^-4^, then we concluded a positive interaction (synergy) was inferred between the two drugs. The functional form for synergy used has also been known as response additivity [44], which was implemented in the Edwards-D-HS model by assuming that the total drug-action is the sum of the individual drug-actions in the absence of any interaction between drugs. The highest single agent (HSA) [45] synergy model is not appropriate here because it does not indicate whether the drugs (AFG with MGX or 5FC) positively interact. It only indicates if their action is higher when administered in combination than maximum drug-action during sole administration. The fractional inhibitory concentration index (FICI) [46] is based on static MIC calculations not conducted here and the Bliss independence model [47] assumes the drugs have independent modes of action, which we do not know to be true. Moreover, response additivity is a more stringent measure of synergy than the Bliss independence model for positive drug-action. We also did not trial implementing Loewe additivity [48] or Zero interaction potency (ZIP) [49] synergy models as these require data from more than one concentration of each antifungal drug, which we do not have in this study.

Finally, the Edwards-D-HS (No Synergy) models assume γ_j_ = ε_j_ = *k*_j_ = 0 for *j* ∈ {AFG: MGX, AFG: 5FC}.

### Model inference

We used the Hamiltonian Monte Carlo (HMC)-based No-U-Turn sampler (NUTs) [50] provided by RStan [51] to sample from all models’ posterior distributions. For each model, four chains were run for 2000 iterations with 50% warm-up and the max_tree_depth parameter set to 12. Signs of non-convergence were monitored using the Gelman-Rubin metric *R^* [52], where *R^* > 1.01 was taken as a sign of non-convergence.

### Model comparison

CV stratified by the replicates was conducted to evaluate the models’ performance on OD_600_ data. When evaluating models on only the drug-free data, one OD_600_ replicate of B12663 growing in RPMI was held out as test data in each fold (*k* = 3). When evaluating models on the individual drug data, OD_600_ replicates in each drug-condition, no drug, AFG, MGX, 5FC, were held out at each CV iteration and iterations were repeated until all the replicates had been in the test-set once (*k* = 3). Similarly, when evaluating models’ performance on all *C. auris* data, including drug combinations, OD_600_ replicates were held out in each of the drug-conditions, no drug, AFG, MGX, 5FC, AFG+MGX and AFG+5FC, and the iterations were rerepeated until all replicates had been in the test set once (*k* = 3).

## Contributions

**T.H.** – Conceptualization, Investigation, Data Curation, Formal Analysis, Methodology, Software, Validation, Visualization, Writing – Original Draft Preparation, Writing – Review & Editing. **L.L.H.J.** – Data Curation, Investigation, Writing – Review & Editing. **E.B.** – Conceptualization, Funding Acquisition, Resources, Writing – Review & Editing. **R.J.T.** – Conceptualization, Project Administration, Funding Acquisition, Resources, Supervision, Writing – Review & Editing.

## Data availability statement

All code used for model fitting and plotting is available on a GitHub repository at https://github.com/tah17/Inferring-mechanistic-synergy-OD-data.

## Financial Disclosure Sentence

We acknowledge funding from the MRC Centre for Medical Mycology at the University of Exeter (MR/N006364/2 and MR/V033417/1), the NIHR Exeter Biomedical Research Centre (NIHR203320) and the MRC Doctoral Training Grant MR/P501955/2. E.B. was also funded by the UK Medical Research Council (MRC) project grant MR/Y002164/1. Additional work may have been undertaken by the University of Exeter Biological Services Unit. The funders had no role in study design, data collection and analysis, decision to publish, or preparation of the manuscript.

## Competing interests

None.

## Related manuscripts

None

## Section

Systems Biology

## Supporting information captions

### S1 Text. Supplementary methods

**Figure S1. Predictive and fitting performance of all models during cross validation on OD_600_ data of B12663 *C. auris*.** Mean relative LPD ((maximum LPD)-LPD + 1) (*top*) per OD_600_ testing replicate averaged over the testing replicates for each fold. Mean RMSE (3.s.f) per OD_600_ testing (*middle*) and training (*bottom*) replicate averaged over the testing and training replicates, respectively, for each fold. Mean and standard errors over folds are shown as dots with grey error bars. Models were assessed on drug-free OD_600_ replicate data (*left*) with the monotherapy drug data (*centre*) and with all the *C. auris* data (*right*). Lower values indicate better predictive and fitting performance on testing and training data, respectively.

**Figure S2. Identifiability of drug parameters in the Edwards-D-HS and Gompertz-D-HS models.** (**a**) Drug-action parameters’ γ_j_ (*blue*), ε_j_ (*green*), *k*_j_ (*red*) for *j* ∈ {AFG, MGX, 5FC, AFG: MGX, AFG: 5FC} posterior contraction, *s*, and z-scores, *z*, when inferred from 5 fake data sets (*left to right*) for Edwards-D-HS (*top*) and Gompertz-D-HS (*bottom*) models. Black dotted lines show the ideal (*s*, *z*)-scores of *s* = 1 and *z* = 0 and grey dotted lines indicate the borders of |z| > 3 which should rarely occur in an unbiased model [21]. (**b**) Distances to the ideal score of (*s*, *z*)=(1, 0) for all estimated drug parameters in the Edwards-D-HS (*purple*) and Gompertz-D-HS (*dark blue*) over the 5 fake data sets. (**c**) Empirical combined coverage of the inferred (1-α)%-credible intervals for the γ_j_ (*blue*), ε_j_ (*green*), *k*_j_ (*red*) parameters. The shaded region shows the 95% binomial proportion confidence intervals for the proportion of times the true parameters were contained in the (1- α)%-credible intervals. At (1- α)-levels of 80% and higher, both models achieve coverage close to the ideal (*dashed line*).

**Figure S3. Edwards-D-HS fake data check results**. 5 sets (*left to right*) of posterior distributions (*dark purple*) for the drug-action parameters ({γ_j_, ε_j_, *k*_j_}_j∈{AFG,MGX,5FC,AFG:MGX,AFG:5FC}_) inferred from 5 fake data sets, which were generated by 5 sets of true parameters (*circles*) arbitrarily sampled from the Edwards-D-HS model’s prior (*light purple*). Dots and thin and thick lines represent the medians and 95% and 80% highest density credible intervals, respectively. The true parameters are contained in the 95% posterior intervals 96% of the time, which is approximately 95% as desired.

**Figure S4. Changes in Edwards and Gompertz-based models’ output when perturbing drug-action parameters from zero.** (**a**) Median of Edwards (*top*) and Gompertz (*bottom*) model outputs in drug-free conditions (*black dashed line*) and in-drug conditions simulated by including one of γ (*blue*), ε (*green*) or *k* (*red*) at a value of 10^-p^ for *p* ∈ {1, …, 6} (*left to right*) in the models. (**b**) MAPE of viable fungal growth (*top*) and RMSE of the median of model outputs (*bottom*) between drug-free and in-drug simulations when each of γ (*blue*), ε (*green*) and *k* (*red*) are perturbed from 0 in the Edwards and Gompertz models (*left to right*). Dotted grey lines indicate our chosen threshold, 10^-4^, where for this value and below parameters were deemed not to change model output.

**Figure S5. Gompertz-D-HS model’s inferred parameter values from OD_600_ data.** Inferred parameter values for AFG (*green*), MGX (*orange*) and 5FC (*purple*) individual drug-action and AFG+MGX (*pink*) and AFG+5FC drug synergy through inhibiting the growth rate (γ.), reduction of the carrying capacity (ε.), and direct killing (*k*.) (*left to right*). Dots and thin and thick lines indicate the medians and the 95% and 80% highest density credible intervals, respectively. Parameter values where inferred 95% credible intervals excluded 10^-4^ were considered to be non-zero and those that included 10^-4^ are shaded grey.

## Notes

https://github.com/tah17/Inferring-mechanistic-synergy-OD-data

### Competing Interest Statement

The authors have declared no competing interest.

## References

[1] Du H, Bing J, Hu T, Ennis CL, Nobile CJ, Huang G. *Candida auris*: Epidemiology, biology, antifungal resistance, and virulence. PLoS Pathog 2020;16:e1008921. 10.1371/JOURNAL.PPAT.1008921.

[2] Lockhart SR, Etienne KA, Vallabhaneni S, Farooqi J, Chowdhary A, Govender NP, et al. Simultaneous emergence of multidrug-resistant *candida auris* on 3 continents confirmed by whole-genome sequencing and epidemiological analyses. Clinical Infectious Diseases 2017;64:134–40. 10.1093/cid/ciw691.

[3] Ostrowsky B, Greenko J, Adams E, Quinn M, O’Brien B, Chaturvedi V, et al. Candida auris Isolates Resistant to Three Classes of Antifungal Medications — New York, 2019. MMWR Morb Mortal Wkly Rep 2020;69:6–9. 10.15585/mmwr.mm6901a2.

[4] World Health Organization. WHO fungal priority pathogens list to guide research, development and public health action. 2022.

[5] CDC. Antibiotic Resistance Threats in the United States. Atlanta, GA: U.S: 2019.

[6] Wake RM, Allebone-Salt PE, John LLH, Caswall BA, Govender NP, Ben-Ami R, et al. Optimizing the Treatment of Invasive Candidiasis - A Case for Combination Therapy. Open Forum Infect Dis 2024;11. 10.1093/ofid/ofae072.

[7] Jaggavarapu S, Burd EM, Weiss DS. Micafungin and amphotericin B synergy against *Candida auris*. Lancet Microbe 2020;1:e314–5. 10.1016/S2666-5247(20)30194-4.

[8] O’Brien B, Liang J, Chaturvedi S, Jacobs JL, Chaturvedi V. Pan-resistant *Candida auris*: New York subcluster susceptible to antifungal combinations. Lancet Microbe 2020;1:e193–4. 10.1016/S2666-5247(20)30090-2.

[9] John LLH, Thomson DD, Bicanic T, Hoenigl M, Brown AJP, Harrison TS, et al. Heightened Efficacy of Anidulafungin When Used in Combination with Manogepix or 5-Flucytosine against *Candida auris In Vitro*. Antimicrob Agents Chemother 2023;67. 10.1128/aac.01645-22.

[10] Spitzer M, Robbins N, Wright GD. Combinatorial strategies for combating invasive fungal infections. Virulence 2017;8:169–85. 10.1080/21505594.2016.1196300.

[11] Bidaud AL, Schwarz P, Herbreteau G, Dannaoui E. Techniques for the assessment of *in vitro* and *in vivo* antifungal combinations. Journal of Fungi 2021;7:1–16. 10.3390/jof7020113.

[12] Aldejohann AM, Wiese-Posselt M, Gastmeier P, Kurzai O. Expert recommendations for prevention and management of *Candida auris* transmission. Mycoses 2022;65:590–8. 10.1111/myc.13445.

[13] Jones CR, Neill C, Borman AM, Budd EL, Cummins M, Fry C, et al. The laboratory investigation, management, and infection prevention and control of *Candida auris*: a narrative review to inform the 2024 national guidance update in England. J Med Microbiol 2024;73. 10.1099/jmm.0.001820.

[14] Kordalewska M, Lee A, Park S, Berrio I, Chowdhary A, Zhao Y, et al. Understanding Echinocandin Resistance in the Emerging Pathogen *Candida auris*. Antimicrob Agents Chemother 2018;62. 10.1128/AAC.00238-18.

[15] Hameed T, Motsi N, Bignell E, Tanaka RJ. Inferring fungal growth rates from optical density data. PLoS Comput Biol 2024;20. 10.1371/journal.pcbi.1012105.

[16] Verhulst P-F. Notice sur la loi que la population suit dans son accroissement. Correspondance Mathématique et Physique Publiée par A. Quetelet 1838;10:113–21.

[17] Gompertz B. On the nature of the function expressive of the law of human mortality, and on a new mode of determining the value of life contingencies. In a letter to Francis Baily, Esq. F. R. S. &c. By Benjamin Gompertz, Esq. F. R. S. Abstracts of the Papers Printed in the Philosophical Transactions of the Royal Society of London 1833;2:252–3. 10.1098/rspl.1815.0271.

[18] Kowalska A, Boruta T, Bizukojc M. Kinetic model to describe the morphological evolution of filamentous fungi during their early stages of growth in the standard submerged and microparticle-enhanced cultivations. Eng Life Sci 2019;19:557–74. 10.1002/elsc.201900013.

[19] Edwards VH. The influence of high substrate concentrations on microbial kinetics. Biotechnol Bioeng 1970;12:679–712. 10.1002/bit.260120504.

[20] Piironen J, Vehtari A. Sparsity information and regularization in the horseshoe and other shrinkage priors. Electron J Stat 2017;11:5018–51. 10.1214/17-EJS1337SI.

[21] Schad DJ, Betancourt M, Vasishth S. Toward a principled Bayesian workflow in cognitive science. Psychol Methods 2021;26:103–26. 10.1037/met0000275.

[22] Dudiuk C, Berrio I, Leonardelli F, Morales-Lopez S, Theill L, Macedo D, et al. Antifungal activity and killing kinetics of anidulafungin, caspofungin and amphotericin B against *Candida auris*. Journal of Antimicrobial Chemotherapy 2019;74:2295–302. 10.1093/jac/dkz178.

[23] Kovács R, Tóth Z, Locke JB, Forgács L, Kardos G, Nagy F, et al. Comparison of In Vitro Killing Activity of Rezafungin, Anidulafungin, Caspofungin, and Micafungin against Four Candida auris Clades in RPMI-1640 in the Absence and Presence of Human Serum. Microorganisms 2021;9:863. 10.3390/microorganisms9040863.

[24] Zhu Y, Kilburn S, Kapoor M, Chaturvedi S, Shaw KJ, Chaturvedi V. *In Vitro* Activity of Manogepix against Multidrug-Resistant and Panresistant *Candida auris* from the New York Outbreak. Antimicrob Agents Chemother 2020;64. 10.1128/AAC.01124-20.

[25] Berkow EL, Lockhart SR. Activity of novel antifungal compound APX001A against a large collection of *Candida auris*. Journal of Antimicrobial Chemotherapy 2018;73:3060–2. 10.1093/jac/dky302.

[26] Pfaller MA, Huband MD, Flamm RK, Bien PA, Castanheira M. Antimicrobial activity of manogepix, a first-in-class antifungal, and comparator agents tested against contemporary invasive fungal isolates from an international surveillance programme (2018-2019). J Glob Antimicrob Resist 2021;26:117–27. 10.1016/j.jgar.2021.04.012.

[27] Watanabe N, Miyazaki M, Horii T, Sagane K, Tsukahara K, Hata K. E1210, a New Broad-Spectrum Antifungal, Suppresses *Candida albicans* Hyphal Growth through Inhibition of Glycosylphosphatidylinositol Biosynthesis. Antimicrob Agents Chemother 2012;56:960–71. 10.1128/AAC.00731-11.

[28] McLellan CA, Whitesell L, King OD, Lancaster AK, Mazitschek R, Lindquist S. Inhibiting GPI anchor biosynthesis in fungi stresses the endoplasmic reticulum and enhances immunogenicity. ACS Chem Biol 2012;7:1520–8. 10.1021/cb300235m.

[29] Liston SD, Whitesell L, Kapoor M, Shaw KJ, Cowen LE. Calcineurin Inhibitors Synergize with Manogepix to Kill Diverse Human Fungal Pathogens. Journal of Fungi 2022;8:1102. 10.3390/jof8101102.

[30] Miyazaki M, Horii T, Hata K, Watanabe N, Nakamoto K, Tanaka K, et al. *In Vitro* Activity of E1210, a Novel Antifungal, against Clinically Important Yeasts and Molds. Antimicrob Agents Chemother 2011;55:4652–8. 10.1128/AAC.00291-11.

[31] Shaw KJ, Ibrahim AS. Fosmanogepix: A Review of the First-in-Class Broad Spectrum Agent for the Treatment of Invasive Fungal Infections. Journal of Fungi 2020;6:239. 10.3390/jof6040239.

[32] Sigera LSM, Denning DW. Flucytosine and its clinical usage. Ther Adv Infect Dis 2023;10. 10.1177/20499361231161387.

[33] Katzir I, Cokol M, Aldridge BB, Alon U. Prediction of ultra-high-order antibiotic combinations based on pairwise interactions. PLoS Comput Biol 2019;15:e1006774. 10.1371/journal.pcbi.1006774.

[34] Greger LM, Greger C, Hiller K-A, Maisch T. Optimal Effective Concentration Combinations and synergy evaluations for binary antimicrobial combinations in vitro. Front Microbiol 2025;16. 10.3389/fmicb.2025.1645341.

[35] Czock D, Keller F. Mechanism-based pharmacokinetic-pharmacodynamic modeling of antimicrobial drug effects. J Pharmacokinet Pharmacodyn 2007;34:727–51. 10.1007/s10928-007-9069-x.

[36] Nielsen EI, Viberg A, Löwdin E, Cars O, Karlsson MO, Sandström M. Semimechanistic pharmacokinetic/pharmacodynamic model for assessment of activity of antibacterial agents from time-kill curve experiments. Antimicrob Agents Chemother 2007;51:128–36. 10.1128/AAC.00604-06.

[37] Tam VH, Schilling AN, Nikolaou M. Modelling time-kill studies to discern the pharmacodynamics of meropenem. Journal of Antimicrobial Chemotherapy 2005;55:699–706. 10.1093/jac/dki086.

[38] Jacobs M, Grégoire N, Couet W, Bulitta JB. Distinguishing Antimicrobial Models with Different Resistance Mechanisms via Population Pharmacodynamic Modeling. PLoS Comput Biol 2016;12. 10.1371/journal.pcbi.1004782.

[39] Pereira LC, de Fátima MA, Santos VV, Brandão CM, Alves IA, Azeredo FJ. Pharmacokinetic/Pharmacodynamic Modeling and Application in Antibacterial and Antifungal Pharmacotherapy: A Narrative Review. Antibiotics 2022;11. 10.3390/antibiotics11080986.

[40] Spalding C, Keen E, Smith DJ, Krachler A-M, Jabbari S. Mathematical modelling of the antibiotic-induced morphological transition of Pseudomonas aeruginosa. PLoS Comput Biol 2018;14:e1006012. 10.1371/journal.pcbi.1006012.

[41] Di Pilato V, Codda G, Ball L, Giacobbe DR, Willison E, Mikulska M, et al. Molecular Epidemiological Investigation of a Nosocomial Cluster of *C. auris*: Evidence of Recent Emergence in Italy and Ease of Transmission during the COVID-19 Pandemic. Journal of Fungi 2021;7:140. 10.3390/jof7020140.

[42] Arendrup MC, Meletiadis J, Mouton JW, Lagrou K, Hamal P, Guinea J, et al. EUCAST Definitive Document E.DEF 7.3.2: Method for the Determination of Broth Dilution Minimum Inhibitory Concentrations of Antifungal Agents for Yeasts. 2020.

[43] Carvalho CM, Polson NG, Scott JG. Handling sparsity via the horseshoe. Journal of Machine Learning Research 2009;5:73–80.

[44] Foucquier J, Guedj M. Analysis of drug combinations: current methodological landscape. Pharmacol Res Perspect 2015;3. 10.1002/prp2.149.

[45] Berenbaum MC. What is synergy? Pharmacol Rev 1989;41:93–141. 10.1016/S0031-6997(25)00026-2.

[46] Fatsis-Kavalopoulos N, Sánchez-Hevia DL, Andersson DI. Beyond the FIC index: the extended information from fractional inhibitory concentrations (FICs). Journal of Antimicrobial Chemotherapy 2024;79:2394–6. 10.1093/jac/dkae233.

[47] Bliss CI. The Toxicity of Poisons Applied Jointly 1. Annals of Applied Biology 1939;26:585–615. 10.1111/j.1744-7348.1939.tb06990.x.

[48] Loewe S. The problem of synergism and antagonism of combined drugs. Arzneimittelforschung 1953;3:285–90.

[49] Yadav B, Wennerberg K, Aittokallio T, Tang J. Searching for Drug Synergy in Complex Dose–Response Landscapes Using an Interaction Potency Model. Comput Struct Biotechnol J 2015;13:504–13. 10.1016/j.csbj.2015.09.001.

[50] Hoffman MD, Gelman A. The No-U-Turn Sampler: Adaptively Setting Path Lengths in Hamiltonian Monte Carlo. Journal of Machine Learning Research 2014;15:1593–623.

[51] Team SD. RStan: the R interface to Stan 2022.

[52] Vehtarh A, Gelman A, Simpson D, Carpenter B, Burkner PC. Rank-Normalization, Folding, and Localization: An Improved R for Assessing Convergence of MCMC. Bayesian Anal 2021;16:667–718. 10.1214/20-BA1221.

