## Supplementary Text for "Inferring antifungal drug synergy from *Candidozyma auris* optical density data using Bayesian mechanistic modelling"

**Supplementary Information**

### A Model details and priors

#### A.1 Edwards

Optical density measured at 600nm ($OD_{600}$) data, $y_t$, measured at a time, $t$, is modelled as a linear transform of true fungal growth, $f(t)$ [cells/$\mu$l], with multiplicative measurement noise at a scale $\sigma$, where the linear transform is parametrised with a proportionality constant, $\frac{1}{\delta}$, and an offset parameter, $B$, representing the average basal $OD_{600}$ [1]. Here $f(t)$ is an Edwards function described using the following ordinary differential equation (ODE):

$$y_t | B, \delta, \sigma, f(0), \beta, L \sim \text{lognormal}\left(\log\left(B + \frac{f(t)}{\delta}\right), \sigma\right),$$
$$\frac{df}{dt} = \beta f(t) e^{-\frac{f(t)}{L}},$$

where $\beta$ is the growth rate and $L$ [cells/$\mu$l] is a constant whose inverse represents the strength of *C. auris*' growth-impedance due to population size. All ODEs are solved by Stan's rk45 ODE solver [2].

The priors of the parameters in this model are:

- $B \sim \text{lognormal}(0, 1)$ for the mean $OD_{600}$ of the background Roswell Park Memorial Institute (RPMI) media. We assume $B$ is always positive and expected to be low in value.
- $\sigma \sim \mathcal{N}^+(0, 0.5)$ for the multiplicative noise scale, where we have used the same prior as previously used when modelling $OD_{600}$ [1] as we expect the $OD_{600}$ reader to have similar measurement noise to other $OD_{600}$ readers. $\mathcal{N}^+$ denotes a normal distribution truncated to the domain $[0, \infty)$ and the distribution $\mathcal{N}^+(0, 0.5)$ is written using its mean, $\mu = 0$, and scale, $\sigma = 0.5$.
- $\tilde{\delta} \sim \mathcal{C}\left(\log_{10}(2.5 \times 10^2), 1\right)$ for the transform of the proportionality constant, $\delta = 10^{\tilde{\delta}}$. Here $\mathcal{C}$ is the Cauchy distribution that is truncated to the domain $[\log_{10}(2.5 \times 10^2), \infty)$. We assume that the median for $\tilde{\delta}$ is $\log_{10}(2.5 \times 10^2)$, corresponding to a $\delta$ value of $2.5 \times 10^2$, which is the initial inoculum size [cells/$\mu$l], and that $\delta > 2.5 \times 10^2$ because wells inoculated with $2.5 \times 10^2$ [cells/$\mu$l] have $OD_{600}$ values that are indistinguishable from $OD_{600}$ of the blanks at time 0.
- $\beta \sim \mathcal{N}^+(0, 1)$ for the growth rate [$h^{-1}$]. We know the growth rate is positive and expect this value to be similar to fungal growth rates previously estimated from $OD_{600}$ [1].
- $\tilde{L} \sim \mathcal{N}(\log_{10}(2.5 \times 10^2), 2)$ for the transform of the inverse strength of *C. auris*' growth-impedance [cells/$\mu$l], $L = 10^{\tilde{L}}$. This assumes that at the initial time, $t = 0$, the growth-limiting term in the ODE, $e^{-\frac{f(0)}{L}}$, is mainly between $e^{-10^2} \approx 0$ and $e^{-10^{-2}} = 0.99$ (3.s.f) $\approx 1$ when $f(0) = 2.5 \times 10^2$.
- $f(0) \sim \text{lognormal}(\log(2.5 \times 10^2), 1)$ for the initial amount of *C. auris* in the well [cells/$\mu$l]. This is assumed to be distributed around the known initial inoculum used in the experiment $2.5 \times 10^2$ [cells/$\mu$l].

For the wells with RMPI only (blanks), the $OD_{600}$ data, $y_{t,b}$, is modelled using $y_{t,b} | \sigma, B \sim \text{lognormal}(\log(B), \sigma)$.

#### A.2 Gompertz

The model has the same likelihood as the Edwards model (Section A.1), but with $f(t)$ being a Gompertz function: $f(t) = f(0)\exp\left\{\log\left(\frac{K}{f(0)}\right)\left(1 - e^{-\beta t}\right)\right\}$, where $\beta$ is now the initial growth rate and is given the same prior as in the Edwards model. All parameters that match with the Edwards model ($B$, $\sigma$, $\tilde{\delta}$, and $f(0)$) have the same definition and priors.

The new parameter, $K$ [cells/$\mu$l], is the carrying capacity. For this parameter we place a prior on the transform of $K$, $\widetilde{K} = \log_{10}(K)$: $\widetilde{K} \sim \mathcal{N}(9, 2)$, where the domain is truncated to $[\log_{10}(2.5 \times 10^2), \infty)$. Lacking further knowledge, we assume the carrying capacity for *C. auris* in the wells will be similar to other fungi but $K$ cannot be less than the initial inocula used in the experiment, and hence we re-use the prior in [1].

#### A.3 Logistic

Similarly to the Gompertz (Section A.2), this model has the same likelihood as the Edwards (Section A.1), but now $f(t)$ is a Logistic function: $f(t) = K\left(1 + \left(\frac{K - f(0)}{f(0)}\right)e^{-\beta t}\right)^{-1}$. The parameters, $B$, $\sigma$, $\tilde{\delta}$, $\beta$ and $f(0)$, are the same as in the Edwards (Section A.1) and $K$ is the same as in the Gompertz model (section A.2) and are given the same priors.

#### A.4 Exponential

As with the Gompertz (Section A.2) and Logistic (Section A.3) models, this model has the same likelihood as the Edwards (Section A.1), but $f(t)$ is an exponential function, $f(t) = f(0)e^{\beta t}$. All parameters, $B$, $\sigma$, $\tilde{\delta}$, $\beta$, and $f(0)$, are the same parameters as in the Edwards model (section A.1) and so are given same definition and priors.

#### A.5 Edwards (OD direct)

$OD_{600}$ data, $y_t$, at a time, $t$, is modelled as directly centred around fungal growth, $f(t)$, with measurement noise of scale $\sigma$. Here, $f(t)$ is an Edwards ODE model solved at initial condition $f(0)$:

$$y_t|\sigma, f(0), \beta, L \sim \text{lognormal}\left(\log(f(t)), \sigma\right),$$
$$\frac{df}{dt} = \beta f(t)e^{-\frac{f(t)}{L}},$$

where $\beta$ is, again, the growth rate and $\frac{1}{L}$ is a constant that represents the strength of *C. auris*' growth-impedance. For the blanks ($f(0) = 0$) the data is modelled using $y_t|\sigma, B \sim \text{lognormal}\left(\log(B), \sigma\right)$, like in Section A.1.

The priors for the parameters $\sigma$, $\beta$ and $B$ are the same as in Section A.1 but for the below parameters we use the following priors:

- $f(0) \sim \mathcal{N}^+(0, 1)$ for the initial $OD_{600}$ value, which we know to be small and positive.
- $L \sim \text{lognormal}(0, 2)$ for the inverse of the growth-impedance constant. We place this prior on this parameter to ensure it is positive and has a domain that allows for $e^{-\frac{f(t)}{L}}$ to be close to 0 and to 1.

#### A.6 Gompertz (OD direct)

Similarly to the Edwards (OD Direct) model (Section A.5), $OD_{600}$ data, $y_t$, at a time, $t$, is modelled as centred around fungal growth, $f(t)$, with measurement noise at scale $\sigma$, but, here, $f(t)$ is the Gompertz function:

$$y_t|\sigma, f(0), \beta, L \sim \text{lognormal}\left(\log(f(t)), \sigma\right),$$
$$f(t) = f(0)\exp\left\{\log\left(\frac{K}{f(0)}\right)\left(1 - e^{-\beta t}\right)\right\},$$

where $\beta$ is the initial growth rate and $K$ is the carrying capacity. Again, for the blanks ($f(0) = 0$) the data is modelled using $y_t|\sigma, B \sim \text{lognormal}\left(\log(B), \sigma\right)$, like in Section A.1.

The priors for $\sigma$, $\beta$, $B$ and $f(0)$ are the same as in the Edwards (OD Direct) model (Section A.5) and the carrying capacity $K$ is given the same prior as $L$ in the Edwards (OD Direct) model (Section A.5): $K \sim \text{lognormal}(0, 2)$, as $OD_{600}$ is known to be positive and generally have maximum values at around 1.

#### A.7 Edwards-indvD

This model extends the Edwards (Section A.1) by including drug-action in the Edwards ODE for the case where there is sole administration of any of the three antifungal drugs: anidulafungin (AFG), manogepix (MGX) and 5-flucytosine (5FC). To include both fungistatic and fungicidal drug-action, we included antifungal drug-action on the growth dynamics and a killing term in the ODEs. This was implemented by including drug-action in the ODE in three ways. First, we assumed that the drugs could inhibit the growth rate, $\beta$, at strengths $\gamma_j$ for each drug $j$. Second, we assumed that the drugs could enhance *C. auris*' growth-impedance, $\frac{1}{L}$, at strengths $\epsilon_j$. Third, and finally, we assumed that each of the drugs could directly kill *C. auris* at a rate $\kappa_j$ [hour$^{-1}$]. Since OD readers cannot differentiate between viable, $f_{\text{v}}(t)$, and dead fungi, $f_{\text{d}}(t)$, the latent *C. auris* growth function, $f(t)$, becomes a sum of the viable and dead *C. auris*, $f(t) = f_{\text{v}} + f_{\text{d}}$:

$$
\begin{aligned}
y_{t,i}|B,\delta,\sigma,f_i(t) &\sim \text{lognormal}\left(\log\left(B + \frac{f_i(t)}{\delta}\right),\sigma\right)\\
f_i(t) &= f_{i,\text{v}}(t) + f_{i,\text{d}}(t),\\
\text{where } \frac{df_{i,\text{v}}}{dt} &= \frac{\beta}{1+\sum_j x_j^i\gamma_j} f_{i,\text{v}}(t) e^{-\frac{f_{i,\text{v}}(t)(1+\sum_j x_j^i \epsilon_j)}{L}} - \left(\sum_j x_j^i \kappa_j\right) f_{i,\text{v}}(t)\\
\frac{df_{i,\text{d}}}{dt} &= \left(\sum_j x_j^i \kappa_j\right) f_{i,\text{v}}(t),
\end{aligned}
$$

for each drug condition $i$ in the wells with *C. auris*; $i \in$ {No drug (RMPI only), AFG, MGX, 5FC}, and each drug $j \in$ {AFG, MGX, 5FC}, where $x_j^i \in \{0,1\}$ is an indicator for the drug $j$'s presence during drug condition $i$. The parameters $B$, $\delta$, $\sigma$, $f(0)$, $\beta$ and $L$ also appear in the Edwards model (Section A.1) and so are given the same prior.

Such that all the priors for the drug-action parameters, $\gamma_j$, $\epsilon_j$ and $\kappa_j$, are on the same scale, the ODEs above are transformed to be dimensionless in time. We did this by scaling time in the ODEs by the maximum observed time, $t_{\text{max}}$, such that the new scaled time $t^\star = \frac{t}{t_{\text{max}}}$ lies in $[0,1]$. As a consequence, we can define a new killing parameter $\kappa_j^\star = t_{\text{max}}\kappa_j$ that is unitless, like $\gamma_j$ and $\epsilon_j$:

$$
\begin{aligned}
\frac{d\tilde{f}_{i,\text{v}}}{dt^\star} &= \frac{t_{\text{max}}\beta}{1+\sum_j x_j^i\gamma_j} \tilde{f}_{i,\text{v}}(t^\star) e^{-\frac{\tilde{f}_{i,\text{v}}(t^\star)(1+\sum_j x_j^i \epsilon_j)}{L}} - \left(\sum_j x_j^i \kappa_j^\star\right) \tilde{f}_{i,\text{v}}(t^\star)\\
\frac{d\tilde{f}_{i,\text{d}}}{dt^\star} &= \left(\sum_j x_j^i \kappa_j^\star\right) \tilde{f}_{i,\text{v}}(t^\star),
\end{aligned}
$$

where $\tilde{f}_{i,\text{v}}(t^\star)$ is $f_{i,\text{v}}(t_{\text{max}}t^\star)$ and the output for the solved ODEs $\tilde{f}_{i,\cdot}(t^\star)$ for $t^\star \in [0,1]$ is the same as the solved ODEs $f_{i,\cdot}(t)$ for $t \in [0, t_{\text{max}}]$. Hence, we solved $\tilde{f}_{i,\cdot}(t^\star)$ for $t^\star \in [0,1]$ and the parameters $\gamma_j$, $\epsilon_j$ and $\kappa_j^\star$ are all given the same standard $\mathcal{N}^+(0,1)$ priors, where the parameters are restricted to be positive in this model (drugs can only act to reduce fungal growth).

For the Edwards-indvD ($\gamma = 0$), Edwards-indvD ($\epsilon = 0$) and Edwards-indvD ($\kappa = 0$), all the priors are kept the same as in the Edwards-indvD model described above, but with $\gamma_j$, $\epsilon_j$ and $\kappa_j$ (and hence $\kappa_j^\star$) set to 0 $\forall j$, respectively.

#### A.8 Gompertz-indvD

This model extends the Gompertz model (Section A.2) by including drug action in the Gompertz ODE for the case where there is sole administration of AFG, MGX or 5FC similarly to the Edwards-indvD model (Section A.7):

$$
\begin{aligned}
y_{t,i}|B,\delta,\sigma,f_i(t) &\sim \text{lognormal}\left(\log\left(B+\frac{f_i(t)}{\delta}\right),\sigma\right)\\
f_i(t) &= f_{i,\text{v}}(t)+f_{i,\text{d}}(t),\\
\text{where } \frac{df_{i,\text{v}}}{dt} &= \frac{\beta}{1+\sum_j x_j^i\gamma_j} f_{i,\text{v}}(t)\log\left(\frac{K}{f_{i,\text{v}}(t)(1+\sum_j x_j^i\epsilon_j)}\right)-\left(\sum_j x_j^i\kappa_j\right)f_{i,\text{v}}(t)\\
\frac{df_{i,\text{d}}}{dt} &= \left(\sum_j x_j^i\kappa_j\right)f_{i,\text{v}}(t),
\end{aligned}
$$

for each drug condition $i$ in the wells with *C. auris* (no drug (RMPI only), AFG, MGX and 5FC) and each drug $j \in \{\text{AFG, MGX, 5FC}\}$, where $x_j^i \in \{0,1\}$ is an indicator for the drug $j$'s presence during drug condition $i$. The ODE for $\frac{df_{i,\text{v}}}{dt}$ can be solved to be:

$$
f_{i,\text{v}}(t) = f_{i,\text{v}}(0)\exp\left\{\left(\log\left(\frac{K}{f_{i,\text{v}}(0)(1+\sum_j x_j^i\epsilon_j)}\right)-\frac{(1+\sum_j x_j^i\gamma_j)\sum_j x_j^i\kappa_j}{\beta}\right)\left(1-\exp\left\{-\left(\frac{\beta}{(1+\sum_j x_j^i\gamma_j)}\right)t\right\}\right)\right\}
$$

The equations are also transformed to be dimensionless in time as in the Edwards-indvD model (Section A.7):

$$
\begin{aligned}
\tilde{f}_{i,\text{v}}(t^\star) &= \tilde{f}_{i,\text{v}}(0)\exp\left\{\left(\log\left(\frac{K}{\tilde{f}_{i,\text{v}}(0)(1+\sum_j x_j^i\epsilon_j)}\right)-\frac{(1+\sum_j x_j^i\gamma_j)\sum_j x_j^i\kappa_j^\star}{t_{\text{max}}\beta}\right)\left(1-\exp\left\{-\left(\frac{t_{\text{max}}\beta}{(1+\sum_j x_j^i\gamma_j)}\right)t^\star\right\}\right)\right\}\\
\frac{d\tilde{f}_{i,\text{d}}}{dt^\star} &= \left(\sum_j x_j^i\kappa_j^\star\right)\tilde{f}_{i,\text{v}}(t^\star),
\end{aligned}
$$

where $t^\star = \frac{t}{t_{\text{max}}}$, $\kappa_j^\star = t_{\text{max}}\kappa_j$ and the $f_{i,\text{v}}(t_{\text{max}}t^\star)$ is rewritten as a new function $\tilde{f}_{i,\text{v}}(t^\star)$.

All the priors for the parameters in this model apart from $K$ are kept the same as in the Edwards-indvD model (Section A.7) and the prior for $K$ is the same as the prior in the Gompertz model (Section A.2).

#### A.9 Edwards-indvD (Decay)

This model extends the Edwards-indvD model (Section A.7) by including decay of dead *C. auris* in the well, $-\delta_\text{d} f_{i,\text{d}}$, to the Edwards ODE of dead fungi, $\frac{df_{i,\text{d}}}{dt}$, for all experimental drug conditions, $i$. All the parameters in this model are kept the same as in the Edwards-indvD model (Section A.7) and the ODEs are also scaled by time to produce $\frac{d\tilde{f}_{i,\text{d}}}{dt^\star}$ with the term $-\delta_\text{d}^\star \tilde{f}_{i,\text{d}}$, where $\delta_\text{d}^\star = t_{\text{max}}\delta_\text{d}$. The new parameter $\delta_\text{d}^\star$ is given a standard $\mathcal{N}^+(0,1)$ prior and is restricted to be positive in this model since it is a decay rate.

#### A.10 Edwards-indvD-HS

The model is the same as the Edwards-indvD model (Section A.7) but with regularised horseshoe priors on the drug-action parameters to enforce sparsity ($\gamma$, $\epsilon$ and $\kappa^\star$) [3]:

$$
\begin{aligned}
\theta &= \{\gamma_j,\epsilon_j,\kappa_j^\star\}_{j\in\{\text{AFG, MGX, 5FC}\}}\\
\theta_k &\sim \mathcal{N}(0,\tilde{\lambda}_k\tau),\ \tilde{\lambda}_k^2 = \frac{c^2\lambda_k^2}{c^2+\tau^2\lambda_k^2}\\
\lambda_k &\sim \mathcal{C}^+(0,1)\\
\tau &\sim \mathcal{C}^+(0,\texttt{scale\_global}\times\sigma)\\
c &= \texttt{slab\_scale}\sqrt{c_{\text{aux}}}\\
c_{\text{aux}} &\sim \text{InvGamma}(0.5\times\texttt{slab\_df},0.5\times\texttt{slab\_df})
\end{aligned}
$$

where we use values of `scale_global` = 1, `slab_scale` = 2 and `slab_df` = 4, which are the default values used in the R package `brms` [4] for the regularised horseshoe. We place the regularising prior on $\gamma$, $\epsilon$ and $\kappa^\star$ as opposed to $\gamma$, $\epsilon$ and $\kappa$ so that all the parameters are regularised on the same scale. Finally, $\gamma$, $\epsilon$ and $\kappa^\star$ are still bounded to be $> 0$ in this model.

##### A.11 Edwards-D-HS (No Synergy)

This model is identical to Edwards-indvD-HS (Section A.10) with identical priors. However, now the model is fit to $OD_{600}$ data with the following 6 drug conditions, $i$, for wells with *C. auris*: no drug (RMPI only), sole administration of AFG, MGX and 5FC and the two drug combinations of AFG+MGX and AFG+5FC.

##### A.12 Gompertz-D-HS (No Synergy)

This model is identical to Gompertz-indvD (Section A.8) with identical priors barring the drug-action parameters, $\gamma_j$, $\epsilon_j$ and $\kappa_j^\star$ that are given the same priors as the drug-action parameters in the Edwards-indvD-HS model (Section A.10). Similarly to the Edwards-D-HS (No Synergy) (Section A.11), the model is now fit to $OD_{600}$ data with all drug conditions, $i$, for wells with *C. auris*: no drug (RMPI only), AFG, MGX, 5FC, AFG+MGX and AFG+5FC.

##### A.13 Edwards-D-HS

This model extends the Edwards-indvD-HS model (Section A.10) to also fit to the $OD_{600}$ data of the combination drug action (AFG+MGX and AFG+5FC). The model has the same equations as the Edwards-indvD-HS model (Section A.10) but now $i$ indexes each drug condition in the wells with *C. auris* (no drug (RMPI only), AFG, MGX, 5FC, AFG+MGX and AFG+5FC) and $j$ now indexes {AFG, MGX, 5FC, AFG:MGX, AFG:5FC}, where AFG:MGX and AFG:5FC are the indices for the interaction parameters. For this model, $x_j^i \in \{0, 1\}$ is now an indicator for if a drug or drug-interaction, $j$, is present during drug condition $i$. For example, if the drug condition $i$ was AFG+MGX, then $x_j^{\text{AFG+MGX}} = 1$ for $j \in$ {AFG, MGX, AFG:MGX} and 0 otherwise. All priors in this model were kept the same as in the Edwards-indvD-HS model (Section A.10).

##### A.14 Gompertz-D-HS

This model extends the Gompertz-D-HS (No Synergy) model (Section A.12) to include synergy terms when fitting to the combination drug $OD_{600}$ data. The model has the same equations as the Gompertz-D-HS (No Synergy) (Section A.12) but now $j$ in the drug-action parameters indexes {AFG, MGX, 5FC, AFG:MGX, AFG:5FC}, where AFG:MGX and AFG:5FC are the indices for the interaction parameters. Similarly to the Edwards-D-HS (Section A.13), $x_j^i \in \{0, 1\}$ in the ODEs is now an indicator for if a drug $j \in$ {AFG, MGX, 5FC} or drug interaction $j \in$ {AFG:MGX, AFG:5FC}, is present during drug condition $i$. All priors in this model were kept the same as in the Gompertz-D-HS (No Synergy) model (Section A.12).

#### B Fake data checks for parameter identifiability

To assess the drug-parameters' identifiability for both Edwards-D-HS (Section A.13) and Gompertz-D-HS (Section A.14), we performed repeated fake data checks [5]. We arbitrarily sampled 5 sets of true parameters, $\{\gamma_j^{\text{true}}, \epsilon_j^{\text{true}}, \kappa_j^{\text{true}}\}$ for $j \in$ {AFG, MGX, 5FC, AFG:MGX, AFG:5FC} from each of the Edwards-D-HS and Gompertz-D-HS models' priors and simulated the corresponding model outputs. These fake data sets were then used to infer 5 sets of drug parameters, $\{\gamma_j, \epsilon_j, \kappa_j\}$, for each model. Ideally, the inferred posteriors should recover their true parameters and have credible intervals that include the true parameters.

To assess how well the true parameters, $\theta^{\text{true}} = \{\gamma_j^{\text{true}}, \epsilon_j^{\text{true}}, \kappa_j^{\text{true}}\}$, were recovered in each of our posterior estimates of the drug-action parameters, $\theta = \{\gamma_j, \epsilon_j, \kappa_j\}$, we calculated the parameters' posterior contraction $s = 1 - \frac{\sigma^2_{\text{post}}}{\sigma^2_{\text{prior}}}$ and the parameters' z-score, $z = \frac{\mu(\theta) - \theta^{\text{true}}}{\sigma_{\text{post}}}$, where $\mu(.)$ is the mean, $\sigma^2_{\text{post}}$ and $\sigma^2_{\text{prior}}$ are the parameters' posterior and prior variance, respectively [6] (Figure **S2a**). For both the Edwards-D-HS and Gompertz-D-HS models, all of the calculated $z$-scores satisfied $|z| < 3$, where $z$-scores with absolute values over 3-4 indicate issues with model identifiability and should rarely occur in unbiased models [6]. Moreover, the majority of both models' $s$-scores were visually near the ideal value of 1 and neither model had a clear visual difference in its distribution of $s$-scores (Figure **S2a**).

To quantitatively compare the identifiability of the drug-action parameters further, we calculated the Euclidean distance from their ($s$, $z$)-scores to the ideal score of (1, 0) for all drug-action parameters inferred from the 5 fake data sets. The models' drug parameters have similar distances to (1, 0) and the densities of these the distances overlap (Figure **S2b**).

The identifiability of the Edwards-D-HS and Gompertz-D-HS models can also be assessed by analysing the empirical coverage of the (1-$\alpha$)%-credible intervals for the drug-action parameters inferred during the fake data check. For a given (1-$\alpha$)-level, coverage was calculated for $\gamma$, for example, by calculating the fraction of times any of true drug-action parameters ($\{\gamma_j^{\text{true}}\}$) were contained in their corresponding estimated parameters' (1-$\alpha$)%-credible interval over all the 5 fake data sets. Coverage was calculated for $\epsilon$ and $\kappa$ in the same way.

Ideally, the (1-$\alpha$)%-credible intervals for the estimated parameters should include the known true parameters (1-$\alpha$)% of the time. If the estimated coverage is above this ideal then the credible intervals are inefficient and they are inaccurate if below.

Both the Edwards-D-HS and Gompertz-D-HS models achieve coverage close to the ideal for $\gamma$, $\epsilon$ and $\kappa$ at (1-$\alpha$)-levels of 80% and higher (Figure **S2c**), where 80% and 95% credible intervals are used for the results in the manuscript. Moreover, the Edwards-D-HS model achieves close to ideal coverage for $\gamma$, $\epsilon$ and $\kappa$ at (1-$\alpha$)-levels of even 50% and higher (Figure **S2c**). In addition to quantitatively calculating the coverage for the Edwards-D-HS model (Figure **S2c**, *left*), we also visually confirmed that the posteriors of the drug parameters inferred from the 5 fake data sets, $\{\gamma_j, \epsilon_j, \kappa_j\}$, contained the true parameters, $\{\gamma_j^{\text{true}}, \epsilon_j^{\text{true}}, \kappa_j^{\text{true}}\}$, that were used to generate the fake data (Figure **S3**).

### C Motivation for choosing $10^{-4}$ as a threshold value for drug-action parameters

To identify which drug-action parameters were indistinguishable from zero, we assessed which inferred drug-action parameters had credible intervals that included values that made no difference to the Edwards-D-HS (Section A.13) and Gompertz-D-HS (Section A.14) models' output. Using the mode of the parameter values that were estimated from fitting these models to the experimental $OD_{600}$ data, we simulated the median drug-free dynamics:

$$B + \frac{f_{\text{No drug, v}} + f_{\text{No drug, d}}}{\delta},$$

for $f$ in both the Edwards (Section A.1) and Gompertz models (Section A.2), which we have denoted as $f_{\text{No drug, v}}$ and $f_{\text{No drug, d}}$ for the viable and dead fungi, respectively. Drug-action was then included into these models by adding one drug-action parameter at a value of $10^{-p}$ for $p \in \{1, \ldots, 6\}$ at a time and then simulating the output; $B + \frac{f_{\text{Drug, v}} + f_{\text{Drug, d}}}{\delta}$, for each added drug-action parameter. Each drug-action parameter was included in the ODEs in a similar manner as for the Edwards-indvD model (Section A.7): $\gamma$ was added as inhibition of growth, $\frac{\beta}{1+\gamma}$, $\epsilon$ as enhancement of growth-impedance in the Edwards, $\frac{1+\epsilon}{L}$, or reduction of carrying capacity in the Gompertz, $\frac{K}{1+\epsilon}$, and $\kappa$ as a direct killing rate, $\frac{df_{\text{Drug, v}}}{dt} = \cdots - \kappa f_{\text{Drug, v}}$. For $p = 4$ and above, no visible difference was observed in the model outputs for all $\gamma$, $\epsilon$ and $\kappa$ (Figure **S4a**).

We also checked which perturbations from 0 for each drug-action parameter resulted in no difference to the ODE simulations of viable growth in the models. Hence, for each perturbation from 0 for $\gamma$, $\epsilon$ and $\kappa$, we measured the mean average percentage error (MAPE) between $f_{\text{No drug, v}}$ and $f_{\text{Drug, v}}$ to assess what perturbation value resulted in less than 1 and 0.1% differences in simulated viable growth, $f_{\cdot,\text{ v}}$. When each drug-action parameter was perturbed from 0 to $10^{-4}$ the corresponding model simulation for viable growth was less than 0.1% different on average from the drug-free simulation (Figure **S4b**, *top*).

Finally, to ensure that perturbing the drug-action parameter by $10^{-4}$ would not make a sizeable difference to the cross validation (CV) results, we also calculated the root mean squared error (RMSE) between $B + \frac{f_{\text{Drug, v}} + f_{\text{Drug, d}}}{\delta}$ and $B + \frac{f_{\text{No drug, v}} + f_{\text{No drug, d}}}{\delta}$ and checked that a perturbation of $10^{-4}$ resulted in RMSEs between simulated in-drug and drug-free growth that was lower than the lowest RMSE recorded on testing or training data during CV. For a perturbation of $10^{-4}$, the RMSE was an order of magnitude below the lowest RMSE in CV (Figure **S4b**, *bottom*).

### References


1. Hameed T, Motsi N, Bignell E and Tanaka RJ. Inferring fungal growth rates from optical density data. PLOS Comput Biol. 2024 May 14;20(5):e1012105.

2. Stan Development Team. 2026. Stan Reference Manual, 2.38. https://mc-stan.org

3. Piironen, J and Vehtari, A. Sparsity information and regularization in the horseshoe and other shrinkage priors. Electron. J. Statist. 2017;11(2):5018-5051

4. Bürkner P. C. brms: An R Package for Bayesian Multilevel Models using Stan. Journal of Statistical Software. 2017;80(1):1-28. doi.org/10.18637/jss.v080.i01

5. Gelman A, Vehtari A, Simpson D, Margossian CC, Carpenter B, Yao Y, Kennedy L, Gabry J, Bürkner PC, Modrák M. Bayesian workflow. arXiv preprint arXiv:2011.01808. 2020 Nov 3.

6. Schad DJ, Betancourt M, and Vasishth S. Toward a principled Bayesian workflow in cognitive science. Psychological methods. 2021 Feb;26(1):103.
