## Supplementary figures and images for "Inferring antifungal drug synergy from *Candidozyma auris* optical density data using Bayesian mechanistic modelling"

### Figure S1

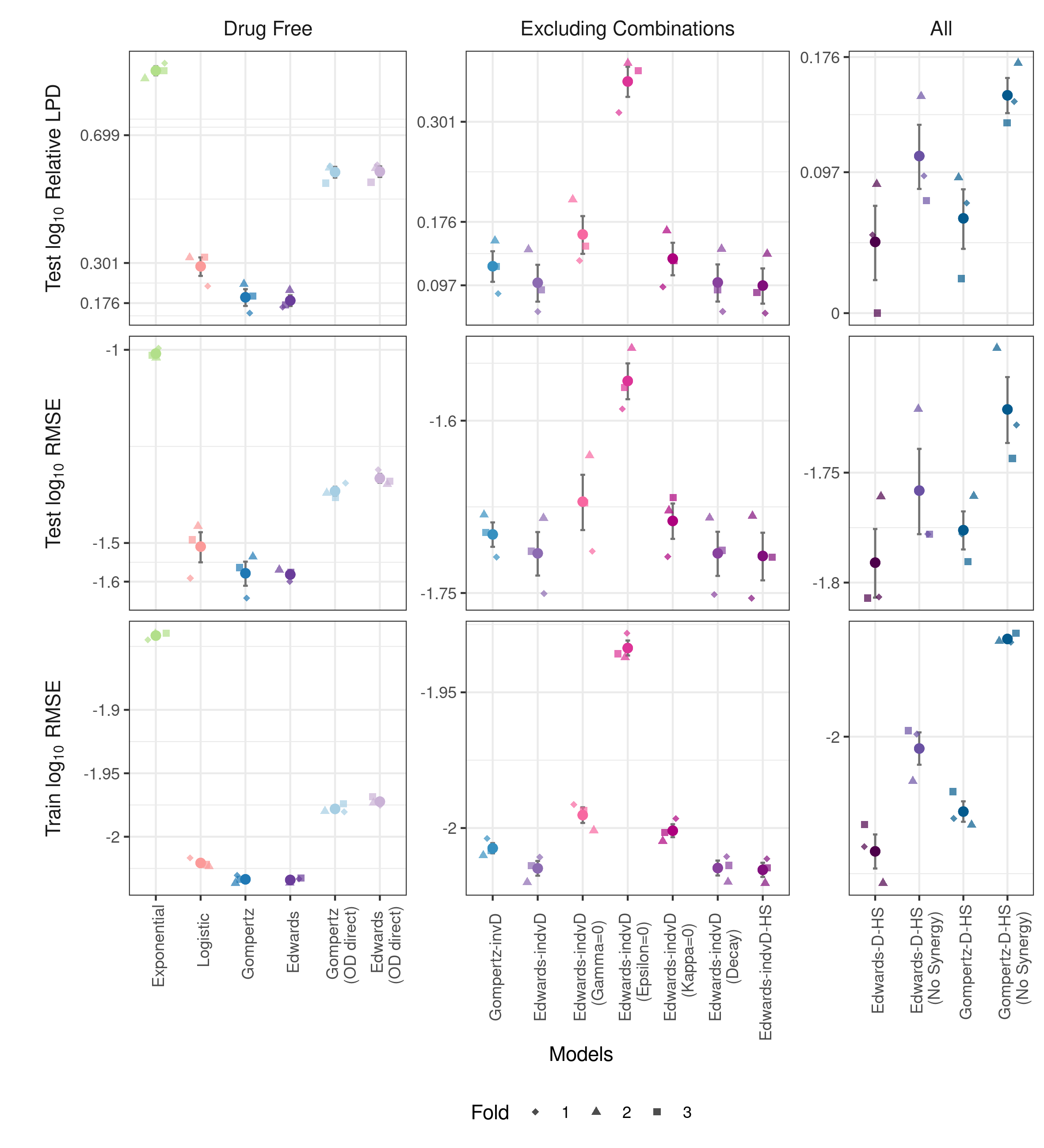

### Figure S2

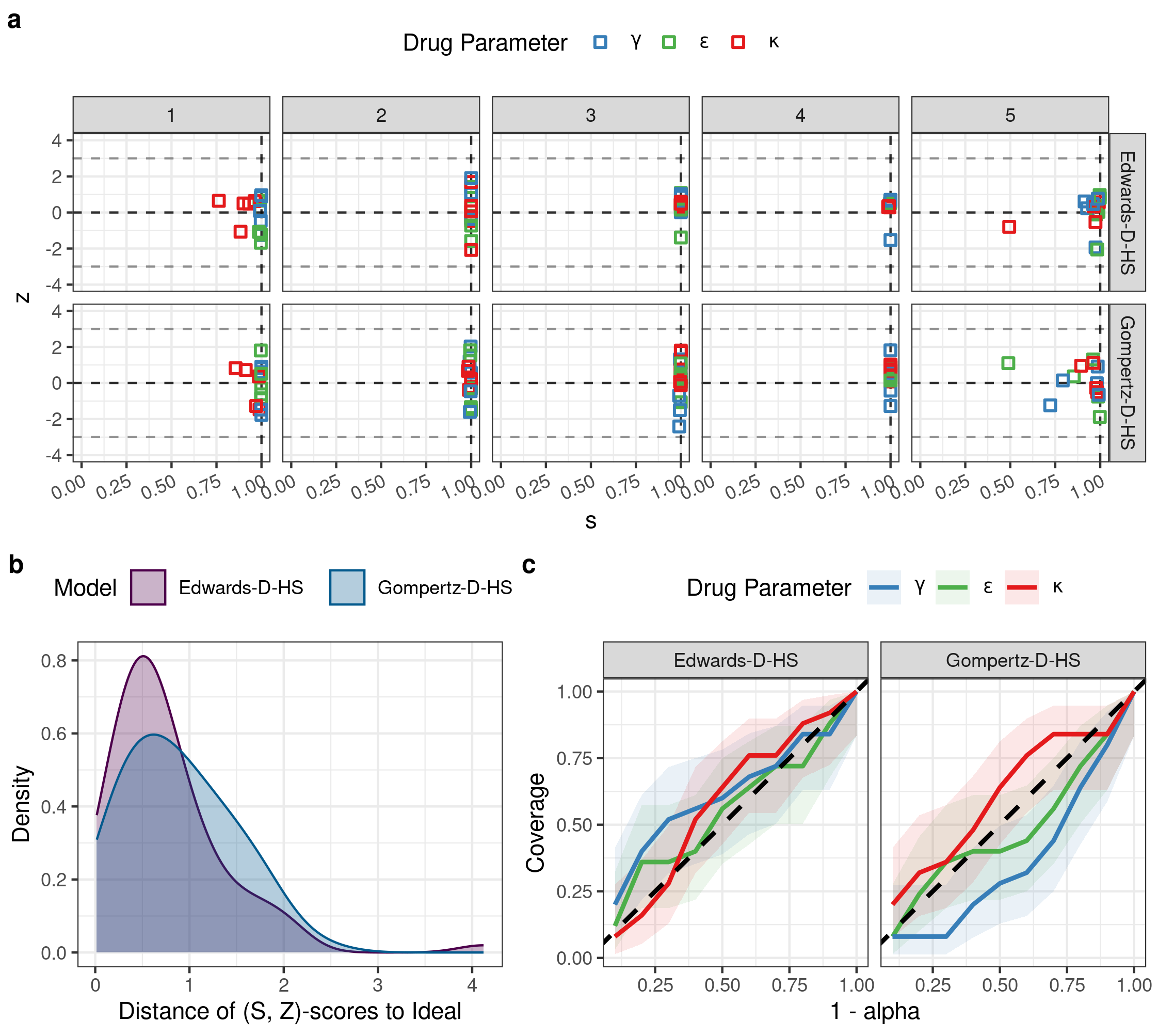

### Figure S3

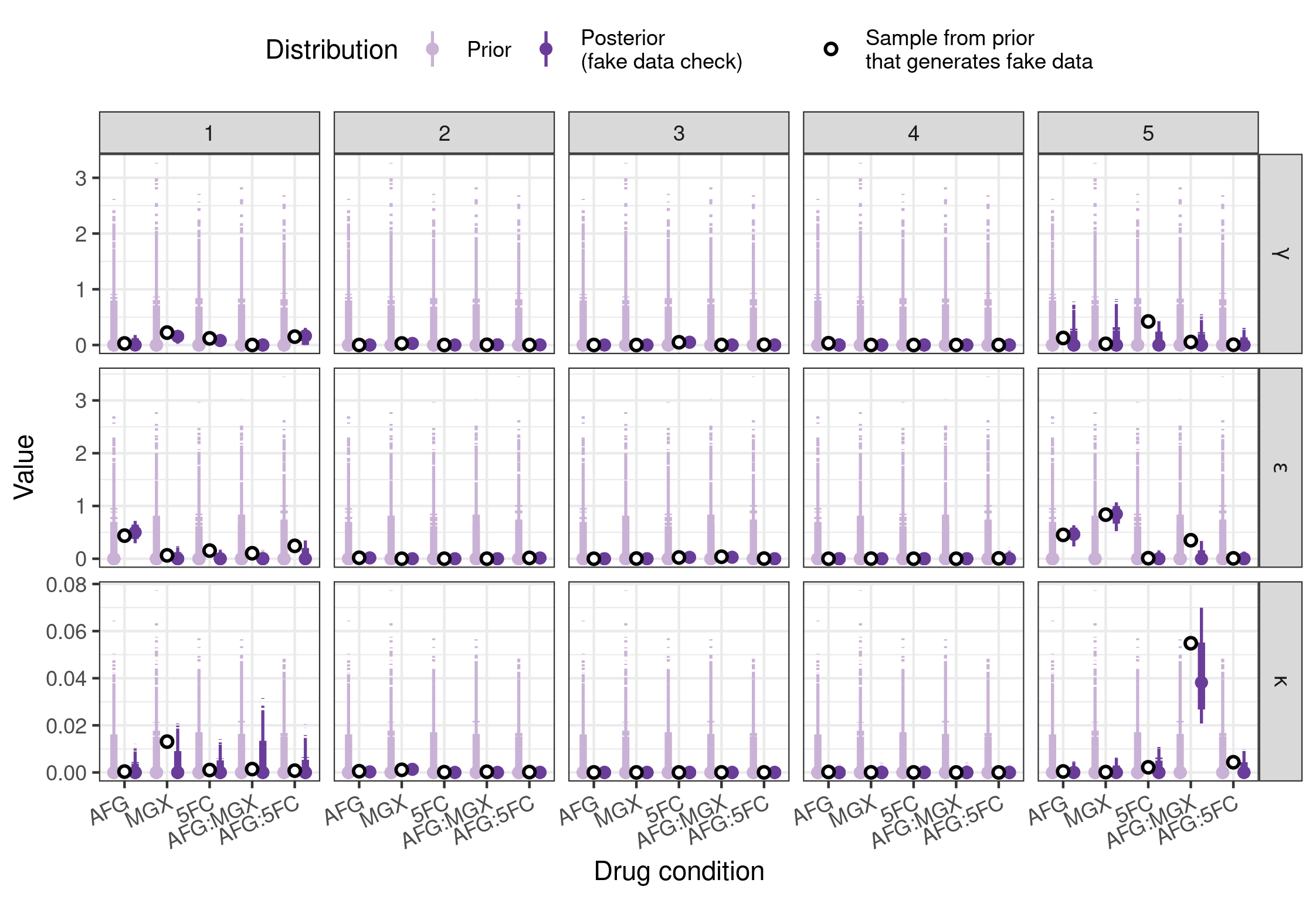

### Figure S4

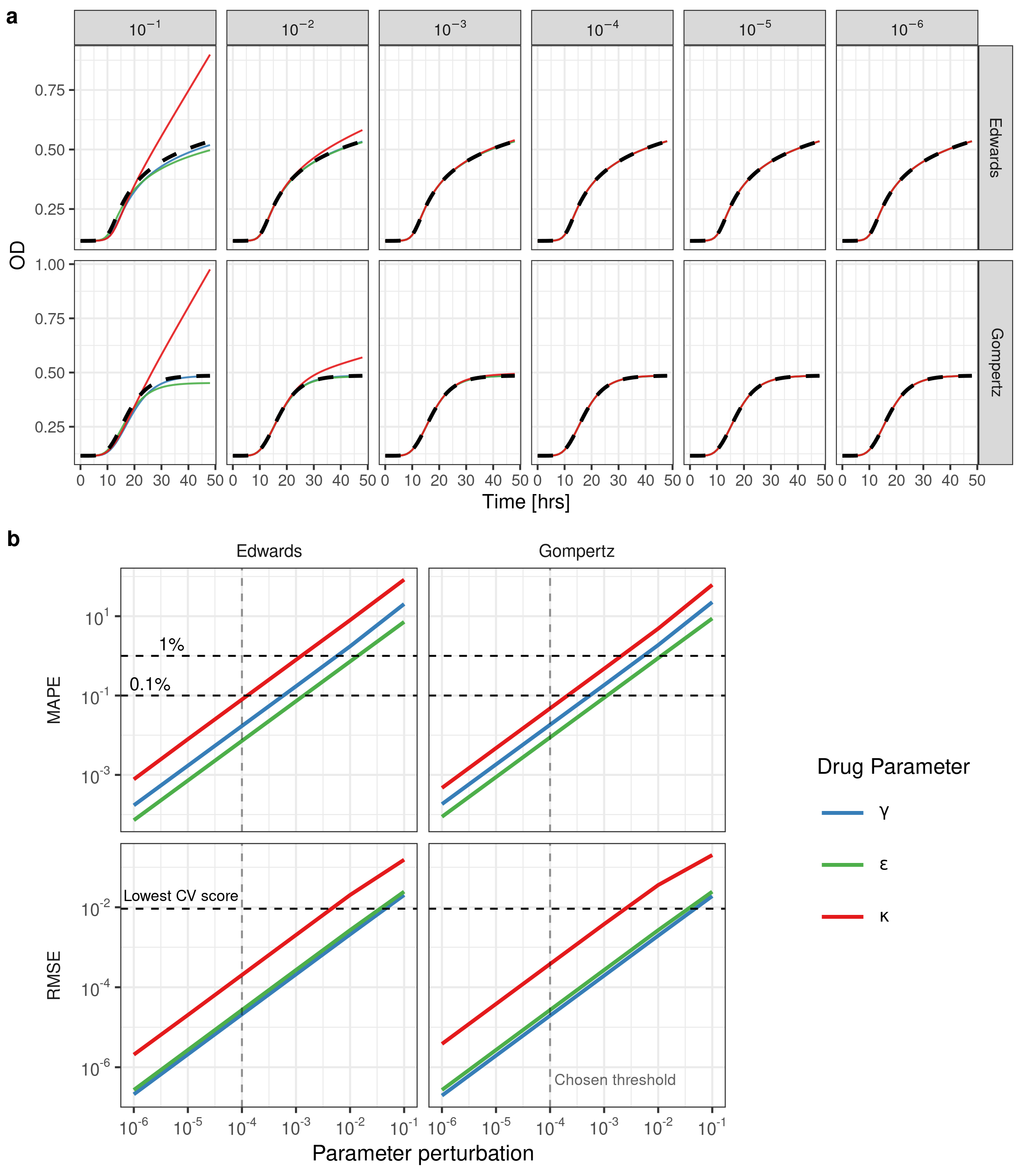

### Figure S5

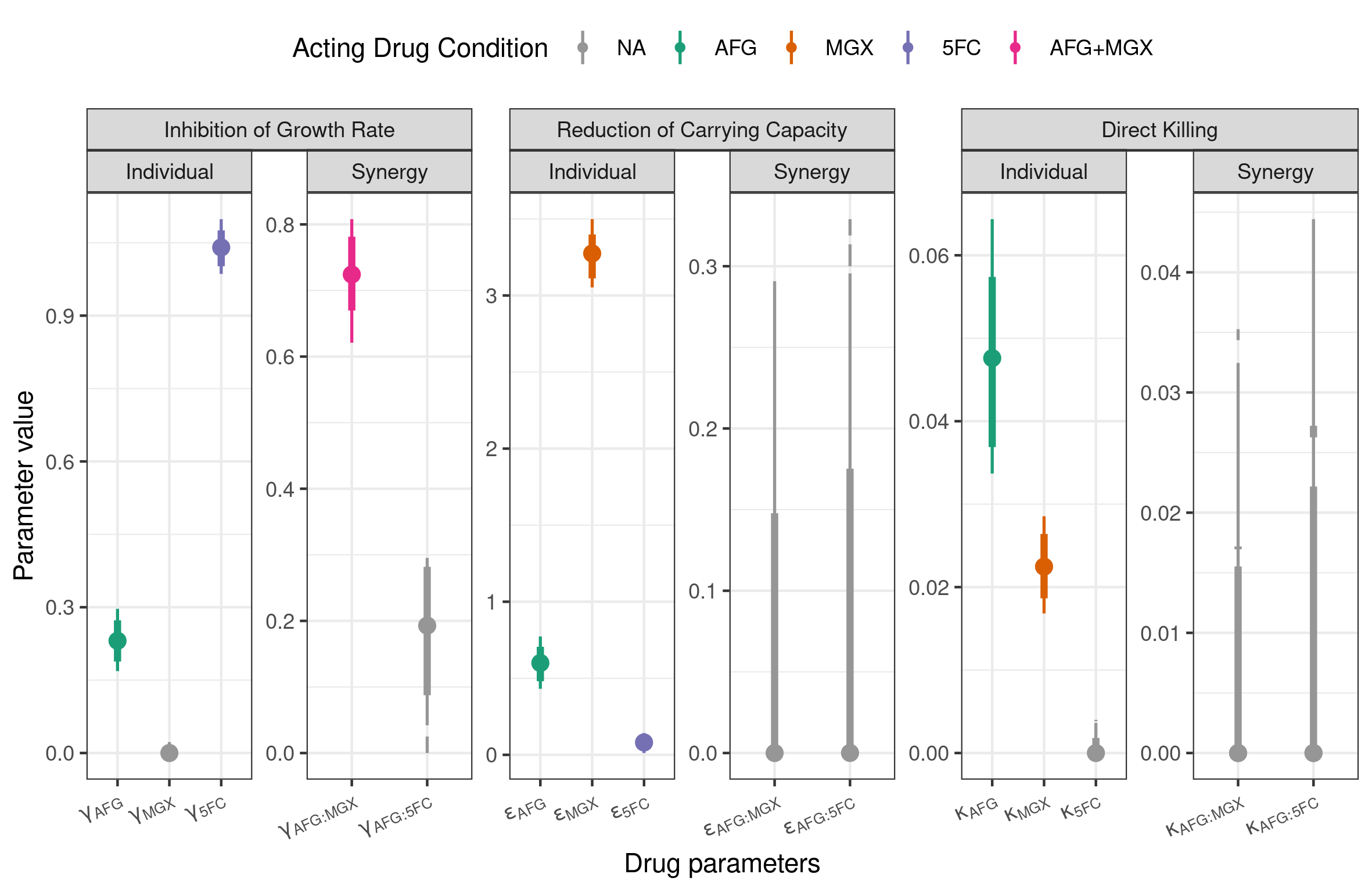
